# Osteocyte Transcriptome Mapping Identifies a Molecular Landscape Controlling Skeletal Homeostasis and Susceptibility to Skeletal Disease

**DOI:** 10.1101/2020.04.20.051409

**Authors:** Scott E. Youlten, John P. Kemp, John G. Logan, Elena J. Ghirardello, Claudio M. Sergio, Michael R. G. Dack, Siobhan E. Guilfoyle, Victoria D. Leitch, Natalie C. Butterfield, Davide Komla-Ebri, Ryan C. Chai, Alexander P. Corr, James T. Smith, Sindhu Mohanty, John A. Morris, Michelle M. McDonald, Julian M. W. Quinn, Amelia R. McGlade, Nenad Bartonicek, Matt Jansson, Konstantinos Hatzikotoulas, Melita D. Irving, Ana Beleza-Meireles, Fernando Rivadeneira, Emma Duncan, J. Brent Richards, David J. Adams, Christopher J. Lelliott, Robert Brink, Tri Giang Phan, John A. Eisman, David M. Evans, Eleftheria Zeggini, Paul A. Baldock, J. H. Duncan Bassett, Graham R. Williams, Peter I. Croucher

## Abstract

Osteocytes are master regulators of the skeleton. We mapped the transcriptome of osteocytes from different skeletal sites, across age and sexes in mice to reveal genes and molecular programs that control this complex cellular-network. We define an *osteocyte transcriptome signature* of 1239 genes that distinguishes osteocytes from other cells. 77% have no previously known role in the skeleton and are enriched for genes regulating neuronal network formation, suggesting this program is important in osteocyte communication. We evaluated 19 skeletal parameters in 733 knockout mouse lines and reveal 26 *osteocyte transcriptome signature* genes that control bone structure and function. We showed *osteocyte transcriptome signature* genes are enriched for human orthologs that cause monogenic skeletal disorders (P=2.4×10^-22^) and are associated with the polygenic diseases osteoporosis (P=1.8×10^-13^) and osteoarthritis (P=1.6×10^-7^). Thus, we reveal the molecular landscape that regulates osteocyte network formation and function and establish the importance of *osteocytes* in human skeletal disease.

## Introduction

The skeleton is a highly dynamic structure that changes in shape and composition throughout life. Osteocytes are the most abundant cell type in bone and have emerged as master regulators of the skeleton. These enigmatic cells are connected via ramifying dendritic processes that form a complex multicellular network distributed throughout mineralized bone^1, 2^. The scale and complexity of the osteocyte network is comparable to neurons in the brain, with 42 billion osteocytes present in the human skeleton forming 23 trillion connections^2, 3^. This network enables osteocytes to detect and respond to mechanical strain, hormones and local growth factors and cytokines^1^. The network responds by regulating the formation and activity of osteoclasts and osteoblasts, instructing these cells to repair damaged bone, controlling bone mass and composition, and ensuring the optimal distribution of bone tissue in response to mechanical stress. Osteocytes also remove and replace bone surrounding the osteocyte network, a surface area of more than 200m^2^ ^3^ by the process of perilacunar remodeling, liberating calcium and phosphate in response to endocrine demands^4^. These features allow the osteocyte network to maintain both the structural integrity of the skeleton and mineral homeostasis. Osteocytes also have regulatory functions beyond the skeleton, including in skeletal muscle, adipose tissue, the central nervous system and in the control of phosphate homeostasis and energy expenditure, indicating the network acts as an important endocrine organ^5–8^.

Although osteocytes are pivotal in controlling the skeleton, the molecular programs that regulate their formation and function are poorly defined. Osteocytes are entombed within bone making them challenging to isolate and study. As a result osteocytes have been omitted from large-scale efforts to map tissue-specific transcriptomes^9–12^ and studies of their transcriptome are limited^13–16^. Consequently, the influences of anatomical location, age and sex on osteocyte regulatory pathways are unclear and their role in the pathogenesis of skeletal diseases is unknown.

Mutations in genes that have highly enriched expression in osteocytes cause rare bone and mineral disorders. For example, autosomal recessive inactivating mutations in *SOST*, which encodes the Wnt-antagonist sclerostin, result in the high bone mass disorder sclerosteosis type 1 (OMIM 269500)^17^. Deletion of a *SOST* regulatory element causes van Buchem disease (OMIM 239100)^18, 19^ and inactivating mutations in *DMP1,* the gene encoding dentin matrix acidic phosphoprotein 1, cause autosomal recessive hypophosphataemia (OMIM 241520)^20^. However, despite the classification of monogenic skeletal disorders identifying over six hundred individual diseases^21^, beyond a limited number of exceptions the role of genes enriched in osteocytes in these disorders remains largely unknown.

Common, complex skeletal diseases, including osteoporosis and osteoarthritis (OA) are also highly heritable. Genetic factors contribute to 50-80% of the variance in bone mineral density (BMD), the major determinant of osteoporosis susceptibility, and account for >50% of the variance in susceptibility to OA^22–24^. Nevertheless, recent large-scale genome-wide association studies (GWAS) have defined only a proportion of the heritability in BMD and OA susceptibility^25, 26^, underscoring the need for alternative approaches to identify genes contributing to these diseases. We hypothesized that mutations in genes with enriched expression in osteocytes would cause rare monogenic skeletal dysplasias and mineral disorders, and contribute to the risk of common polygenic skeletal diseases including osteoporosis and osteoarthritis.

To address this hypothesis, we developed a method to define a transcriptomic map of the osteocyte network and used it to investigate relationships between genes with enriched expression in osteocytes and human skeletal disease. We defined the genes expressed in osteocytes isolated from different skeletal sites, at different ages and from both sexes, and discovered bone-specific and sexually dimorphic differences in the osteocyte transcriptome during post-natal development. We defined an *osteocyte transcriptome signature,* a profile of genes with enriched expression in osteocytes, and discovered novel genes and molecular programs that control formation and function of the osteocyte network. Finally, we showed that *osteocyte transcriptome signature* genes are (i) highly enriched for genes that cause monogenic skeletal disorders and (ii) significantly over-represented with susceptibility genes for BMD and OA identified by human GWAS.

## Results

### The Osteocyte-Enriched Transcriptome is Highly Conserved Throughout the Skeleton

To investigate the genes that control osteocytes, we first identified the repertoire of genes expressed in osteocytes, the osteocyte-enriched transcriptome, from three different skeletal sites (**Fig. 1a-b**). Total RNA was isolated from skeletal samples (98% cortical bone, 2% cancellous bone) highly enriched for osteocytes (>80% of cells present) from the tibiae, femora and humeri of 16-week old mice (n=8) and sequenced (**Fig. 1b-c** and **Supplementary Fig 1a-d**). A threshold of ‘active’ gene expression was determined based on the distribution of normalised gene expression in each sample^27^ (**Fig. 1d, Supplementary Table 1**).

**Fig. 1:**
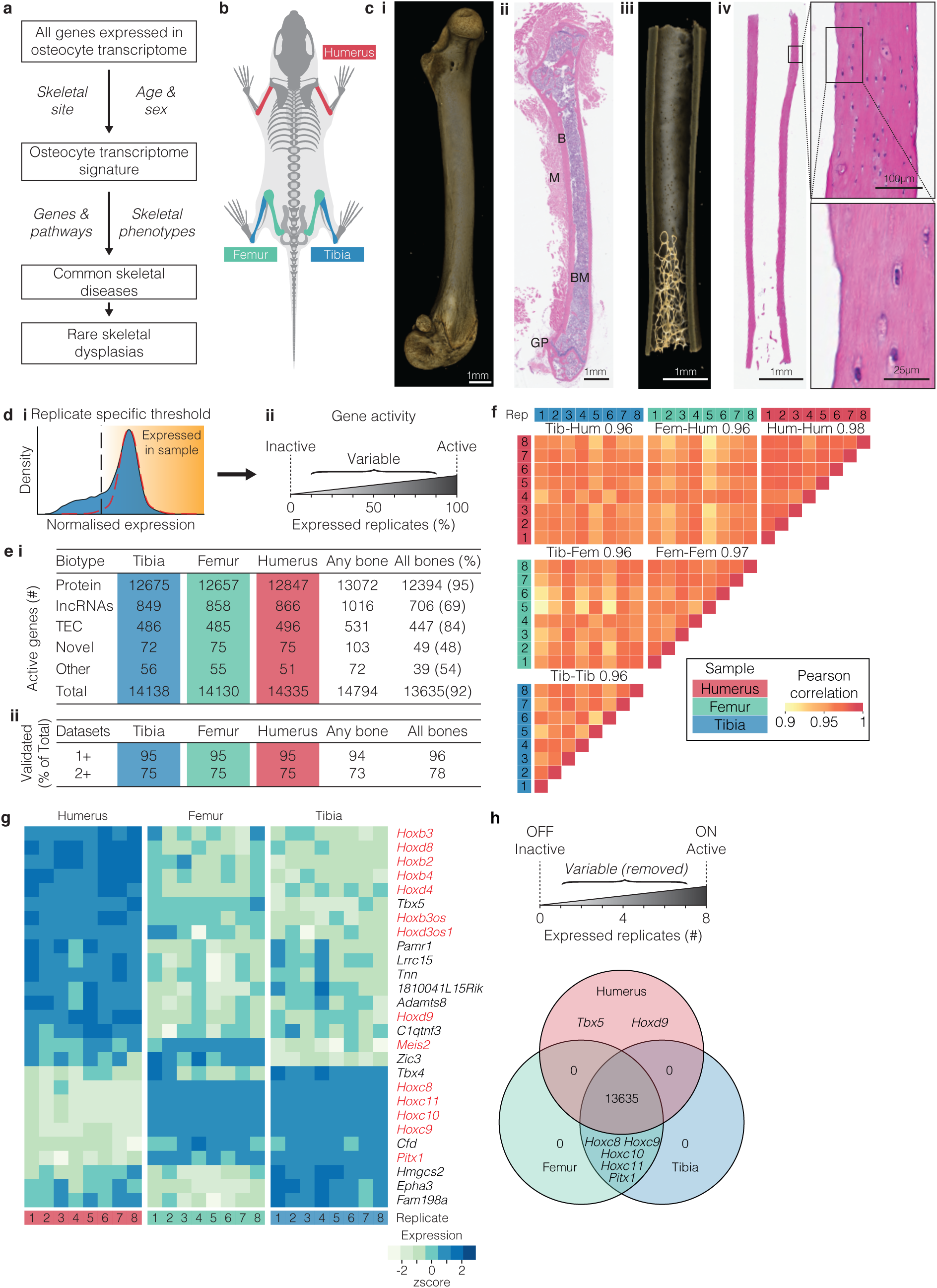
The osteocyte transcriptome is broadly conserved across the skeleton. **a,** Schematic of the overall study design used to define the osteocyte transcriptome, to define an *osteocyte transcriptome* signature, the profile of genes enriched in osteocytes, and to identify their role in skeletal disease. **b,** Diagram illustrating the 3 bone-types in which the osteocyte transcriptome was sequenced and compared. **c,** Representative micro-CT and histological images of bone samples prior to processing **(i-ii)** and following osteocyte isolation **(iii-iv)**, illustrating the effective removal of bone-marrow, muscle and growth-plates and enrichment of osteocytes **(iv).** Boxes and dotted lines identify areas of high magnification confirming osteocyte isolation. Bone (B), bone marrow (BM), muscle (M) and growth-plate (GP) are identified. **d,** Actively expressed genes were identified based on the normalized gene expression distribution (blue histogram) and a hypothetical distribution of ‘active’ promoters (red dotted line)^27^ **(i)**. Vertical line denotes the sample-specific threshold of actively expressed genes. Genes were considered ‘active’ if they were above sample-specific threshold in all replicates, ‘inactive’ if they were below the threshold in all replicates, or ‘variable’ if they were above the threshold in at least one but not all replicates **(ii)**. **e,** The number of ‘active’ genes and biotype composition in the osteocyte transcriptome across skeletal sites **(i)**, and the percentage of total active genes detected in osteocytes in at least one (1+) and two or more (2+) orthogonal datasets **(ii)**. The ‘Any bone’ column reflects the number of genes actively expressed at any skeletal site. The ‘All bones’ column reflects the number and proportion (as a percentage) of genes that were actively expressed in all skeletal sites. TEC = To be Experimentally Confirmed. **f,** Pearson correlation of gene expression between individual replicates and bone types (mean of each bone type comparison represented by numbers above heatmap). Tib = tibia, Fem = femur, Hum = humerus, Rep = biological replicate. **g,** Genes that were differentially expressed between osteocytes isolated from different bone types. Homeobox genes and antisense-RNAs are in red (FDR ≤ 0.05, LFC > 0.5). **h,** Methodology for identifying genes expressed in a site-specific manner and a Venn diagram identifying the genes expressed in one bone type but not others.

14,794 genes were actively expressed in the three bone types with 92% of genes expressed at all sites and 96% of genes validated in one of three independent datasets, including the IDG-SW3 osteocytic cell-line, laser-capture micro-dissected osteocytes or osteocytes isolated by collagenase digestion (**Fig. 1e-f, Supplementary Table 2**). These comprised protein-coding genes, long non-coding RNAs (lncRNAs), genes present in the GENCODE transcriptome annotation but yet to be experimentally confirmed (TEC), and novel genes yet to be reported in any public annotation (**Fig. 1e**). The number of genes actively expressed in osteocytes was similar to other tissues, including 8997 genes found in a dataset of twelve organs and tissues^28^ (**Supplementary Fig. 2a**, **Supplementary Table 2**). Osteocytes formed a separate cluster in a principal component analysis (PCA), confirming the osteocyte-enriched transcriptome is clearly distinct from the transcriptome of other tissues (**Supplementary Fig. 2b**). In support of this, analysis of gene expression specificity, using the Tau measure^29^ calculated across these tissues, showed a significant proportion of genes with high specificity of expression for osteocytes (**Supplementary Fig. 2c**).

Whilst the osteocyte-enriched transcriptome was conserved among bones, 27 genes were differentially expressed between skeletal sites (LFC>0.5, p<0.05, **Fig. 1g, Supplementary Table 3**). These included 7 genes expressed specifically in either the fore (2 genes) or hindlimb (5 genes) (**Fig. 1h**). All encoded developmental transcription factors. They included T-box 5 (*Tbx5*) and homeobox-d9 (*Hoxd9*), expressed exclusively in the humerus and known to establish forelimb identity^30, 31^. By contrast, Homeobox-c8-c11 (*Hoxc8, Hoxc9, Hoxc10* and *Hoxc11*) and paired-like homeodomain 1 (*Pitx1*) were expressed in the femur and tibia only, and not the humerus. In the developing limb bud, *Pitx1* is expressed exclusively in the hindlimb and is the master regulator of hindlimb-type identity^32^. Aberrant expression of *PITX1* in the forelimb leads to homeotic arm-to-leg transformation in Liebenberg syndrome (OMIM 186550)^33^. Fifty-two percent of the 27 differentially expressed genes, were homeobox genes, or *Hox*-antisense lncRNAs, indicating this family may be important in maintaining the identity of osteocytes at different sites (**Fig. 1h**). Together these data show that the osteocyte transcriptome is highly conserved at different anatomical sites, although, homeobox genes, which are typically associated with patterning in development, identify osteocytes from different skeletal locations, even in adult mice.

### Sex and Age Differences in the Osteocyte-Enriched Transcriptome

Since bone structure and bone mass vary with sex and age^34–36^, we hypothesized that the osteocyte-enriched transcriptome differs between the sexes and changes with age. We therefore analysed bone structure and the transcriptome of male and female mice at different ages (**Fig. 2a**). Bone length and bone mineral content (BMC), but not bone mineral density (BMD), differed between sexes, whereas bone length, BMC and BMD increased with age in both sexes (**Fig. 2b, Supplementary Fig. 3)**. The total number of genes actively expressed by osteocytes increased with skeletal maturation with 81% of genes expressed at all ages in both sexes (**Fig. 2c, Supplementary Table 2**). Comparison of gene expression between age groups showed differences between 4 to 10 weeks of age (female p=8×10^-3^, male p=3×10^-5^), but not from 10-16 weeks or 16-26 weeks, demonstrating the osteocyte transcriptome expressed during growth is distinct from the transcriptome in the adult skeleton (**Fig. 2d**). Comparison of the osteocyte transcriptome between sexes showed it was different between male and female mice in the mature skeleton (16 weeks, p=8.0×10^-3^ and 26 weeks, p=2.8×10^-2^), but not at earlier ages (**Fig. 2e**).

**Fig. 2:**
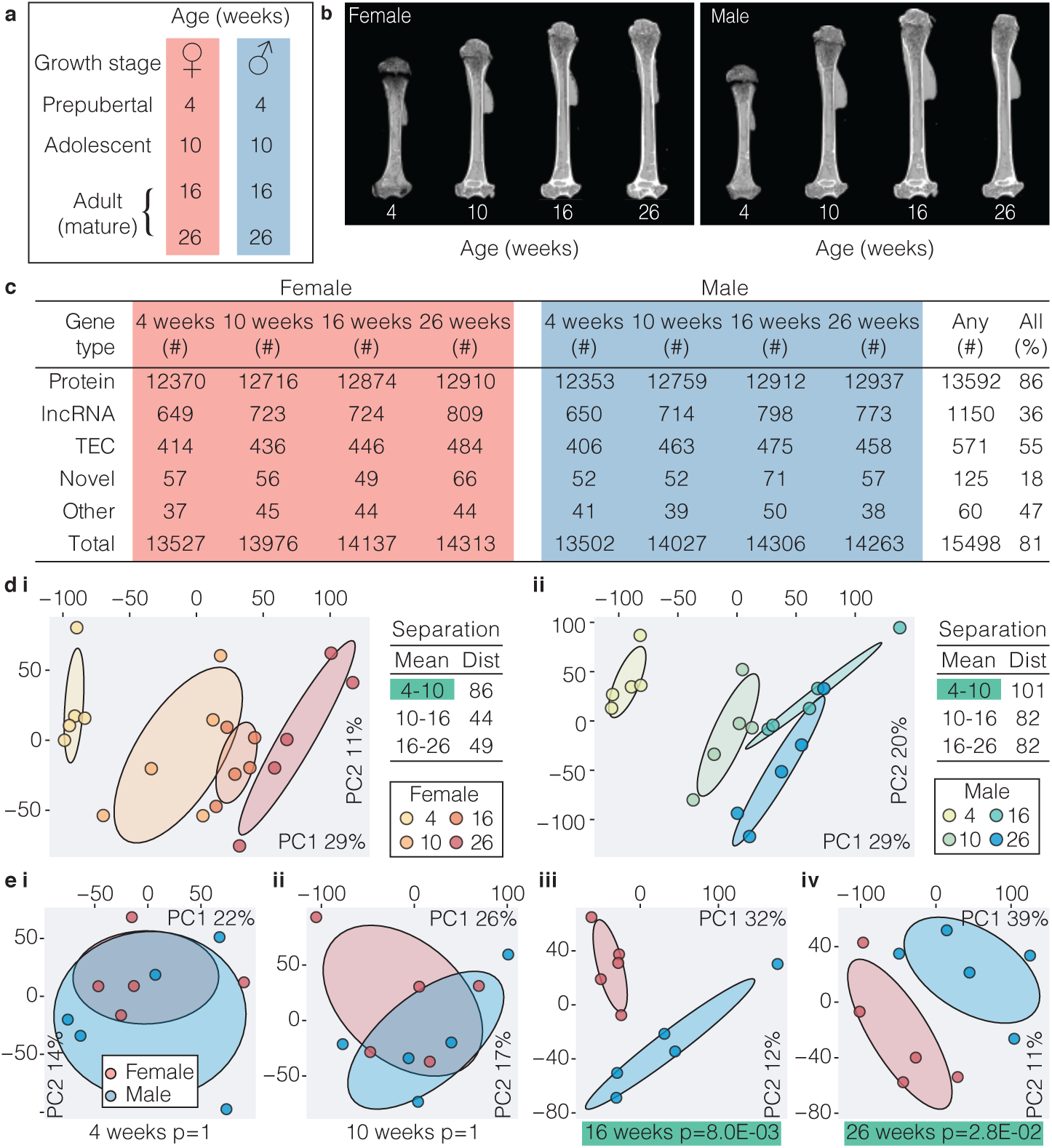
The osteocyte transcriptome changes with sex and age. **a,** Experimental design used to define the osteocyte transcriptome in each sex during skeletal maturation. **b,** Representative micro-CT images of humeri from female and male mice at different ages. **c,** The number of active genes and biotype composition of the osteocyte transcriptome at different ages in each sex. ’Any’ reflects the number of genes actively expressed in all ages and sexes. TEC = to be experimentally confirmed. **d,** Principal component (PC) analysis of samples clustered by age in female **(i)** and male **(ii)** mice. The percentage of total variance explained by individual PCs is shown. Dots represent individual biological replicates and ellipses represent 50% confidence intervals for each age. Dist = Euclidean distance between cluster centroids (mean). Clusters with significant separation between centroids are highlighted in green (p ≤ 0.05). **e,** Principal component analysis of samples clustered from both sexes at 4 **(i)**, 10 **(ii)**, 16 **(iii)** and 26 **(iv)** weeks of age. The percentage of total variance explained by individual PCs is shown. Dots represent individual biological replicates and ellipses represent 50% confidence intervals for each sex. Clusters with significant separation between centroids are highlighted in green (p≤

T o identify genes and processes that contribute to differences in age and sex, clusters of co-expressed genes were identified by weighted gene co-expression network analysis. Seven clusters of correlated genes (denoted by seven colours) were identified (**Supplementary Fig. 4a** and **Supplementary Table 4**). One cluster (denoted by *Grey*) contained genes that were not correlated with each other or other clusters. Between 86% and 97% of genes in each cluster were also found in at least one of three orthogonal datasets, including the IDG-SW3 osteocytic cell line, micro-dissected osteocytes or osteocytes isolated by collagenase digestion (**Supplementary Fig. 4b)**. Each cluster was associated with distinct biological processes (**Supplementary Fig. 4b**). The expression of genes within two clusters (*Purple* and *Turquoise*) increased with age, whereas, gene expression in two clusters (*Black* and *Brown*) decreased with age (**Supplementary Fig. 4a**). The expression of genes in the *Brown* and *Magenta* clusters were associated with both age and sex (**Supplementary Fig. 4a** and **Supplementary Fig. 5a**). The *Brown* cluster contained genes encoding bone matrix constituents, including osteocalcin (*Bglap*, *Bglap2),* osteonectin (*Sparc*) and bone sialoprotein (*Ibsp*) and was associated with protein-processing and transport (**Supplementary Fig. 4a** and **Supplementary Table 4**). The *Magenta* cluster included cathepsin K (*Ctsk*), tartrate resistant acid phosphatase (*Acp5*) and the vacuolar ATPase family and was associated with processes relating to *bone resorption* (GO:0045453, p=4.37×10^-6^), *osteoclast differentiation* (GO:0030316, p=1.40×10^-8^), *pH reduction* (GO:0045851, p=2.31×10^-6^) and *ATP-coupled cation transport* (GO:0099132, p=2.13×10^-14^) (**Supplementary Fig. 5b-c** and **Supplementary Table 4**). To exclude a contribution to the *Magenta* cluster from genes expressed in contaminating osteoclasts we stained paraffin sections of osteocyte-enriched bone samples for tartrate resistant acid phosphatase (TRAP) and confirmed the absence of TRAP positive osteoclasts on trabecular and cortical endosteal bone surfaces (**Supplementary Fig. 5d**). Furthermore, analysis of the top 20 *Magenta* cluster genes, which include genes typically found in osteoclasts, showed that 19 were found in at least one of the three orthogonal datasets (**Supplementary Fig. 5e**). Together this suggests the *Magenta* cluster genes may be important in regulation of perilacunar-resorption^15, 37^. To investigate further, we examined expression of *Magenta* cluster genes in lactating mice in a publicly available microarray dataset^15^. Sixty-six of the 84 *Magenta* genes were up-regulated during lactation (p=3.8×10^-35^) and this was reversed post-lactation (p=1.2×10^-15^) (**Supplementary Fig. 5f**) strengthening the notion that the *Magenta* cluster identifies genes involved in perilacunar remodeling. Together, this highlights the dynamic regulation of the osteocyte transcriptome during post-natal skeletal development and identifies clusters of genes that are differentially regulated during skeletal maturation and between the sexes.

### A Unique Transcriptome Signature Defines the Osteocyte

To identify the genes that distinguish osteocytes specifically, we identified a profile of genes whose expression was enriched in osteocytes relative to other cell types. We hypothesized that genes important for osteocyte-specific functions are actively expressed and preferentially expressed in osteocytes compared to other cell lineages within bone, including bone marrow cells and cells lining bone (**Fig. 3a**). To test this, we performed transcriptome analysis on bone samples enriched with osteocytes, from which bone marrow and cells lining bone were removed, and compared this to whole bone samples, in which bone marrow was retained (**Fig 3a**). The expression of genes encoding established osteocyte proteins were among the most enriched in osteocyte enriched bone samples (**Fig. 3b, Supplementary Table 2**). *Sost* and *Mepe* were enriched by >100-fold and *Dmp1* >40 fold^38^, whereas the expression of house-keeping genes was unaffected by cell composition and genes typically expressed in bone marrow cells were depleted in osteocyte-enriched samples (**Fig. 3b**).

**Fig. 3:**
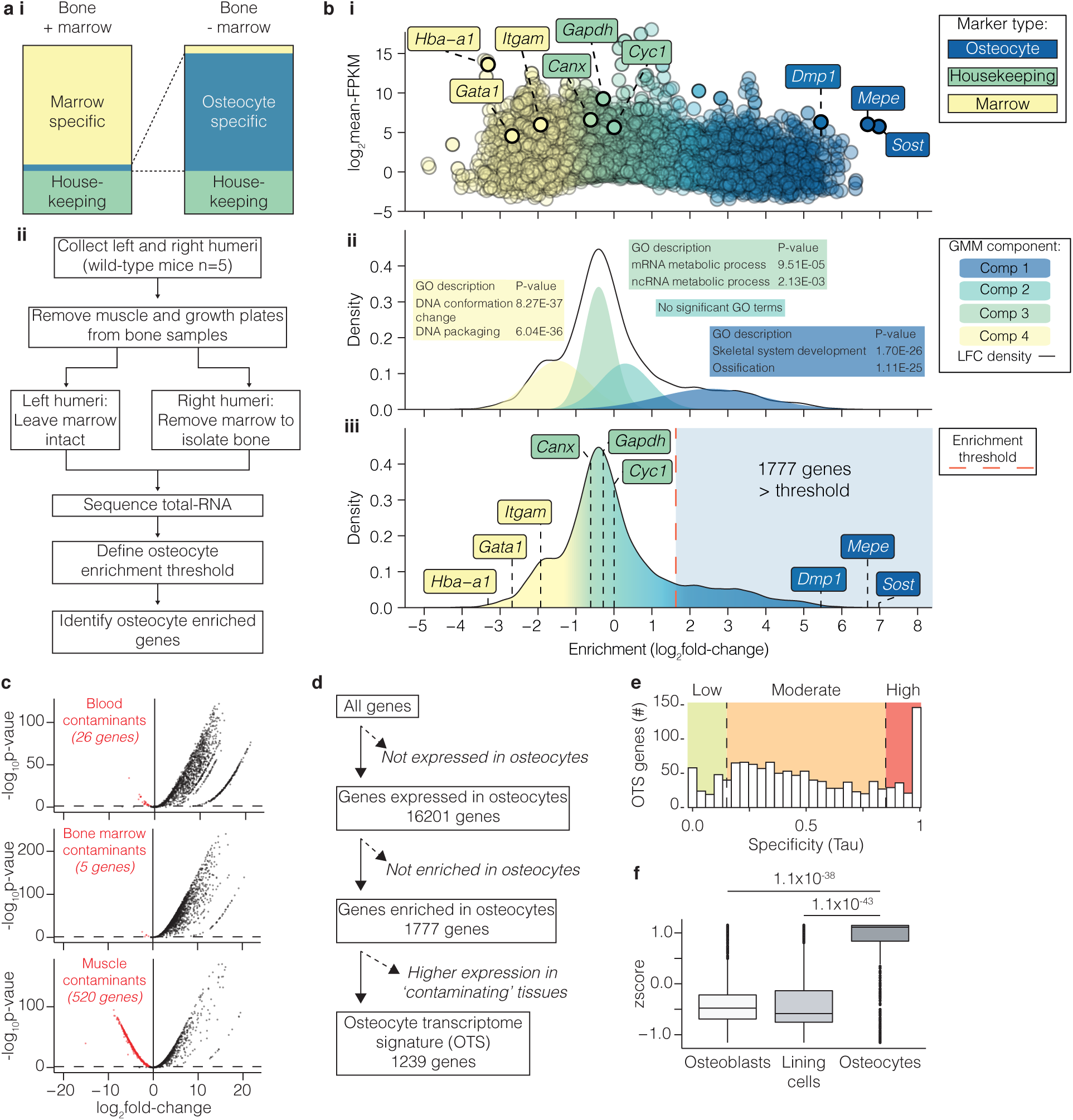
Defining an *osteocyte transcriptome signature*. **a,** Experimental design used to identify genes with enriched expression in osteocytes. Diagram illustrating the strategy used to identify genes enriched for expression in osteocytes in bone samples enriched with osteocytes relative to intact bone samples in which bone marrow and cells lining bone were retained **(i)**. Experimental workflow used to identify genes enriched for expression in osteocytes **(ii)**. **b,** Gene-enrichment in osteocyte-enriched bone samples relative to whole-bone samples distinguished genes known to be expressed in osteocytes (blue) from housekeeping genes (green) and genes expressed in marrow cell populations (yellow) **(i)**. Enrichment for each individual gene is shown as a dot and plotted as a function of normalised gene expression (log_2_-FPKM). The 4 component gaussian mixture model fitted to the density distribution of gene-enrichment in **(i)** used to define the osteocyte enrichment threshold **(ii)**. Each component is denoted by a separate colour. The top two Gene Ontology biological processes associated with genes in each component are illustrated in coloured boxes with p-values (Bonferroni-corrected) **(ii)**. 1777 genes above the osteocyte-enrichment threshold (indicated by the red dashed line) were significantly enriched for expression in osteocytes **(iii). c,** Volcano plots comparing the expression of osteocyte-enriched genes (from **b**) with blood **(i)**, bone-marrow **(ii)** and muscle **(iii)**^13^. Dashed lines represent p<0.05 cutoff. Genes significantly enriched in these tissues relative to osteocytes are identified as red dots. **d,** Filtering pipeline used to define the 1239-gene *osteocyte transcriptome signature* (OTS). The number of genes remaining at each filtering stage is indicated. **e,** Expression specificity^29^ (Tau) of OTS genes relative to other organs and tissues^28^. Genes with Tau < 0.15 = low expression specificity in osteocytes (green), 0.15 ≤ Tau ≤ 0.85 = moderate expression specificity (orange), while Tau > 0.85 = high expression specificity (red). **f,** OTS genes were enriched for expression in osteocytes relative to osteoblasts and bone lining cells isolated by laser capture microdissection. Tukey box-plots show a summary of median OTS gene expression values in each cell type. Boxes indicate median and interquartile range (IQR) of scaled, normalized gene expression values, whiskers denote values ± 1.5*IQR and outlier values beyond this range are shown as individual points. P-values were calculated by CAMERA^105^.

Next, we fitted a 4 component Gaussian Mixture Model (GMM) to the distribution of gene enrichment and used this to calculate a threshold of osteocyte-enrichment. This model identified 1777 genes with significantly enriched expression in osteocytes (**Fig. 3b, Supplementary Table 2**). As an additional level of stringency 538 genes enriched in bone marrow or tissues that could contaminate the osteocyte enrichment strategy, such as blood or muscle (**Fig. 3c, Supplementary Table 2**)^13^, were excluded leaving 1239 genes significantly enriched for expression in osteocytes **(Fig. 3d, Supplementary Table 5**). 85% of these genes showed moderate to high expression specificity^29^, using the Tau measure, within the osteocyte network relative to 12 non-skeletal tissues (**Fig. 3e**). Furthermore, osteocyte enriched genes were highly expressed in osteocytes relative to osteoblasts (P=1.1×10^-38^) and bone lining cells (P=1.1×10^-43^) in a publicly available microarray dataset (**Fig. 3f**)^39^. Using this pipeline we thus defined a list of 1239 genes whose expression is enriched in osteocytes relative to bone marrow and other cells in the osteoblast lineage. We defined this profile of genes as the *osteocyte transcriptome signature* (**Supplementary Table 5, Fig. 3d**).

### The Majority of *Osteocyte Transcriptome Signature* Genes Have No Known Function in the Skeleton

Analysis of the *osteocyte transcriptome signature* showed it was enriched with genes associated with skeletal biological processes in the GO database^40^ (4.5 fold-enrichment (FE), P=1.0×10^-67^), and with skeletal phenotypes in the Mouse Genome Informatics database^41^ (MGI, 2.7 FE, P=4.7×10^-35^). This included *Sost*, *Dkk1*, *Mepe* and *Dmp1*, genes known to be highly expressed in osteocytes, and genes with an established role in the skeleton, such as osteoprotegerin (*Tnfsf11b*), Wingless-type family member-1 (*Wnt1*)^42^, fibroblast growth factor-9 (*Fgf9*)^43^ and Iroquois homeobox protein 5 (*Irx5*)^44^ (**Fig. 4a, Supplementary Fig 6**). Interestingly, *Tnfsf11* encoding RANKL (the ligand for receptor activator of NFkB), was expressed by osteocytes but not present in the *osteocyte transcriptome signature* (**Supplementary Fig 6**). A limited number of genes were not annotated with skeletal terms in the GO database, but have been reported to have a role in the skeleton (denoted as ‘reported’) (**Fig. 4a**). They include the Wnt-regulator notum (*Notum*), which regulates bone formation^45–47^ and a distintegrin and metalloproteinase like member (*Adamtsl2*), which is implicated in geleophysic dysplasia 1 (OMIM 231050)^48^. The majority of *osteocyte transcriptome signature* genes (78%, n=968) had not previously been shown to have a role in the skeleton (‘unannotated’ in **Fig. 4a**).

**Fig. 4:**
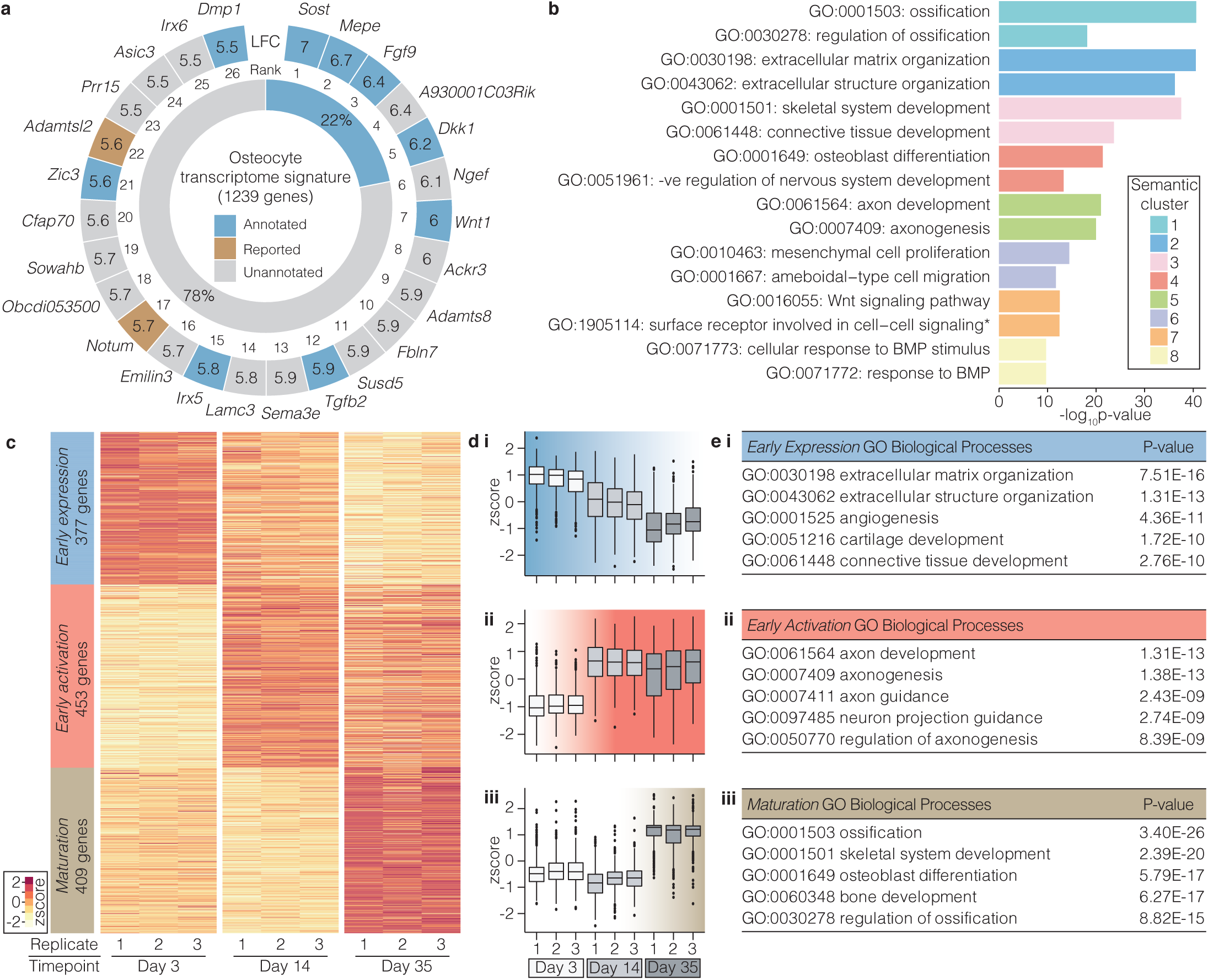
Genes and control pathways identified by the *osteocyte transcriptome signature*. **a,** The top 26 *osteocyte transcriptome signature* genes most enriched in osteocytes (outer ring). Genes ‘annotated’ with either a skeletal biological process (GO database) or skeletal phenotype (MGI database) are highlighted blue; genes ‘reported’ with experimental evidence of a role in the skeleton are brown, whereas ‘unannotated’ genes without a skeletal annotation in GO or MGI and no experimental evidence of a role in the skeleton reported in the literature are shown in grey. Numbers denote log_2_ fold-enrichment (LFC) in gene expression in osteocytes. The proportion of all *osteocyte transcriptome signature* genes ‘annotated’ (blue) or unannotated (grey) with a skeletal annotation in GO or MGI is shown in the inner ring. **b,** Clusters (n=8) of semantically similar biological processes (GO) significantly enriched in the *osteocyte transcriptome signature* are identified by different colours (Bonferroni-corrected p<0.05). The top 2 terms (by p-value) in each of the 8 clusters are listed. **c,** Heatmap showing clustering of *osteocyte transcriptome signature* genes based on distinct co-expression patterns during osteocytic-differentiation^50^ of the IDG-SW3 cell-line, from osteoblast-like cells (day 3) to early (day 14) and mature osteocytes (day 35). The name and number of genes for each cluster are labelled. Cluster colours correspond to subsequent panels **(d-e)**. **d,** Changes in *early expression* cluster **(i)**, *early activation* cluster **(ii)** and *maturation* cluster **(iii)** of genes during osteocyte differentiation. Tukey box-plots show the distribution of gene expression values for each cluster in each replicate. Boxes indicate median and interquartile range (IQR) of scaled, normalized gene expression values, whiskers denote values ± 1.5*IQR and outlier values beyond this range are shown as individual points. **e,** Biological processes (BP) enriched in the *early expression* cluster (i), the *early activation* cluster (ii) and the *maturation* cluster (iii). The top 5 GO BP terms in each cluster and enrichment p values are shown.

In addition to known genes, forty-nine novel genes were actively expressed in osteocytes (**Supplementary Fig. 7a**, **Supplementary Table 6**). Eleven were present in the *osteocyte transcriptome signature,* (**Supplementary Fig. 7b**), including 7 that were absent from 12 other tissues (**Supplementary Fig. 7c**). The multiple exons and splicing patterns suggested post-transcriptional processing of transcripts, while analysis of sequence-coding potential indicated they were all non-coding genes (**Supplementary Fig. 7c**). Thus, the *osteocyte transcriptome signature* expands the repertoire of genes whose expression is enriched in osteocytes and includes known and novel genes.

Analysis of GO terms enriched in the *osteocyte transcriptome signature* identified 8 semantically similar clusters of related processes (**Fig. 4b**, **Supplementary Table 7**). Cluster 1 was enriched with processes associated with ‘*ossification’* (GO:0001503, p=2.3×10^-41^), cluster 2 *‘extracellular matrix organization’* (GO:0030196, p=2.8×10^-41^), cluster 3 ‘*skeletal system development’* (GO:0001501, p=2.9×10^-38^), cluster 4 ‘*osteoblast differentiation’* (GO:0001649, p=4.2×10^-22^) and cluster 6 ‘*mesenchymal cell proliferation*’ (GO:0010463, p= 3.44×10^-15^), whereas, clusters 7 and 8 were enriched with processes associated with signaling, including ‘*wnt signaling pathway’* (GO:0016055, p=3.0×10^-13^) and *‘cellular response to BMP stimulus*’ (GO:0071773, p=2.0×10^-10^, **Fig. 4b**). GO term analysis also identified cluster 5 which was enriched with terms associated with axon guidance including ‘*axon development’* (GO:0061564, p=9.6×10^-22^) and *‘axonogenesis’* (GO:0007409, p=1.0×10^-20^) (**Fig. 4b**). ‘*Axon guidance’* (mmu04360, p=7.8×10^-17^) was also the top ranked KEGG pathway (**Supplementary Fig. 8a**). This included genes in the Semaphorin, Ephrin, Netrin and Slit signaling families and their respective receptors Plexins, Eph-receptors, Uncoordinated-5 (*Unc-5*) and Roundabout (*Robo*), which are pivotal regulators of axonal guidance (**Supplementary Fig. 8b**). Since axon guidance directs the formation of the intercellular neuronal network^49^, we hypothesized that this pathway is a key molecular program required for osteocyte network formation.

To examine this, we investigated temporal patterns of *osteocyte transcriptome signature* gene expression in the *in vitro* IDG-SW3 model of osteocyte cell differentiation in which cells differentiate from late osteoblast-like cells, via early osteocytes, to late osteocytic cells^50^. Three clusters of genes were identified (**Fig 4c**, **Supplementary Table 5**), an *early expression cluster* - 377 genes most highly expressed in osteoblast-like cells, but down-regulated as cells transition to early and mature osteocytes (**Fig. 4c-d**); an *early activation cluster* – 453 genes up-regulated in early osteocytes, and which remained expressed in mature osteocytes (**Fig. 4c-d**); and a *maturation cluster* – 409 genes up-regulated in mature osteocytes (**Fig. 4c-d**). The *early expression cluster* was enriched for processes associated with *‘extracellular matrix organisation’* (GO:0030198, p=7.51×10^-16^, **Fig. 4e**) and the maturation cluster with ‘*ossification’* (GO:0001503, p=3.40×10^-26^, **Fig. 4e**). The *early activation cluster* was associated with *‘axon development’* (GO:0061564, p=1.31×10^-13^), ‘*axonogenesis*’ (GO:0007409, p=1.38×10^-13^) and ‘*axon guidance’* (GO:0007411, p=2.43×10^-9^, **Fig. 4e**). The upregulation of these processes coinciding with early osteocyte differentiation and sustained expression in late osteocytes suggests axonal guidance pathways are important in the formation and maintenance of the osteocyte network.

### *Osteocyte Transcriptome Signature* Genes Control Bone Structure and Function

To establish whether *osteocyte transcriptome signature* genes have a functional role in the skeleton, we screened mice with single gene deletions that have undergone detailed skeletal phenotyping by the *Origin of Bone and Cartilage Disease* (*OBCD*) program (http://www.boneandcartilage.com/bonepipeline.html). Structural and functional bone phenotyping has been performed in 733 knockout mouse lines, of which 64 had deletions of genes present in the *osteocyte transcriptome signature* (**Fig. 5a**, **Supplementary Table 8**). 26 (41%) of these knockout lines had either structural and/or functional skeletal phenotypes. Eleven were in genes with established roles in the skeleton, including *Daam2*^25^ and *Pls3*^51^. Fifteen were in genes not annotated with skeletal terms or phenotypes in the GO or MGI databases, suggesting they are novel regulators of bone structure and/or function (**Fig. 5a, Supplementary Table 8**). These included the activator of transcription and development regulator (*Auts2*), a regulator of neuron migration^52^, dapper antagonist of beta-catenin 3 (*Dact3*), which controls Wnt signaling^53, 54^ and low-density lipoprotein receptor class A domain containing protein (*Ldlrad4*), an inhibitor of TGF-Beta signaling^55^ (**Fig. 5a**). Genes in the MGI database, but without functional evidence of a role in the skeleton such as cortactin binding protein 2 (*Cttnbp2*) (**Fig. 5a**), a regulator of dendrite arbourisation^56^, were also identified. *Auts2^-/-^* mice had decreased femoral bone length and cortical diameter (**Fig. 5b-c**), whereas *Cttnbp2^-/-^* had normal femoral length, BMC and functional parameters but increased vertebral BMC and maximum load (**Fig. 5b,c,e**). *Dact3*^-/-^ had normal femoral bone length, BMC, bone structural parameters and functional parameters, however, BMC was increased in the vertebra and this was accompanied by increased yield load (**Fig. 5b,c,e**). *Ldlrad4^-/-^* had normal bone length, BMC and functional parameters but had decreased bone volume (BV/TV), decreased trabecular number (Tb.N) and increased trabecular separation (Tb.Sp) (**Fig. 5b,c**). High resolution micro-CT analysis of osteocyte lacunae showed that deletion of *Cttnbp2* and *Ldlrad4* caused an increase in lacuna volume with increased numbers of large lacunae and fewer smaller lacunae, deletion of *Auts2* enhanced lacuna sphericity, whereas *Dact3* deletion had no effect on size distribution or sphericity (**Fig. 5d, Supplementary Fig. 9**). *Auts2* and *Ldlrad4* expression was higher in osteocyte-enriched bone than in other tissues. By contrast, expression of *Cttnbp2* and *Dact3* was highest in brain and/or adrenal gland, respectively (**Supplementary Fig. 10**), suggesting that deletion of these two genes might result in both direct local effects on the skeleton and secondary additional contributions via their expression in other tissues.

**Fig. 5:**
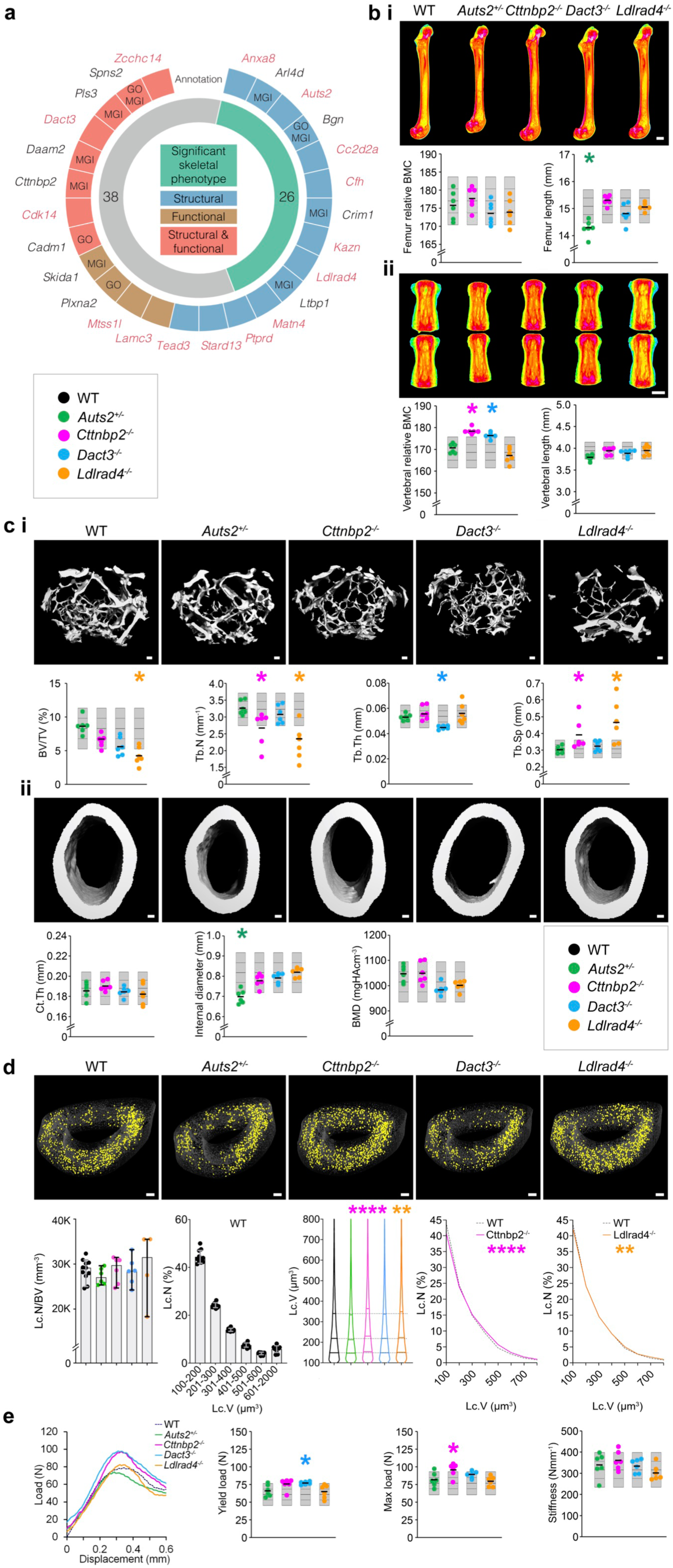
Deletion of *osteocyte transcriptome signature* genes affects bone structure and function. **a,** The 26 *osteocyte transcriptome signature* genes with outlier skeletal phenotypes in the Origins of Bone and Cartilage Disease (OBCD) database of single gene deletions in mice (outer ring). Deletion of genes in blue affected bone structure, genes in brown affected bone function and genes in red both structure and function. Those annotated in either GO or MGI are indicated. Genes in red have a significant outlier phenotype when deleted in knockout mice but are not annotated with skeletal function or phenotype in either GO or MGI databases. The inner ring denotes the number of genes with (green) and without (grey) an outlier skeletal phenotype when deleted in mice in the OBCD database. **b,** Representative quantitative X-ray microradiographic images from the femurs **(i)** and vertebrae **(ii)** of adult, female wild-type (WT), *Auts2*^+/-^, *Cttnbp2*^-/-^, *Dact3*^-/-^ and *Ldlrad4*^-/-^ mice. Scale bar = 1 mm. Dot plots illustrate individual parameters. **c,** Representative micro-CT images of trabecular **(i)** and cortical **(ii)** bone of adult, female wild-type (WT), *Auts2*^+/-^, *Cttnbp2*^-/-^, *Dact3*^-/-^ and *Ldlrad4*^-/-^ mice. Scale L 100µm. Dot plots illustrate bone volume as a proportion of tissue volume (BV/TV), trabecular number (Tb.N), trabecular thickness (Tb.Th), trabecular separation (Tb.Sp), cortical thickness (Ct.Th), internal endosteal diameter and bone mineral density (BMD). **d,** Representative micro-CT images showing large osteocyte lacunae (401-800 µm^3^) in tibia cortical bone from adult, female WT, *Auts2*^+/-^, *Cttnbp2*^-/-^, *Dact3*^-/-^ and *Ldlrad4*^-/-^ mice. Scale bar = 100 μ m. Graphs show osteocyte lacunae number per bone volume (Lc.N/BV) and distribution of lacunae volumes (Lc.V) in WT mice. Violin plot shows distribution of Lc.V in the four knockout mouse lines compared to WT and graphs show relative frequency distribution of Lc.V in *Cttnbp2*^-/-^ and *Ldlrad4*^-/-^ mice compared to WT (n = 4-11 per genotype). ** P<0.01, **** P<0.0001. **e,** Load displacement curves from caudal vertebrae compression testing in adult female WT, *Auts2*^+/-^, *Cttnbp2*^-/-^, *Dact3*^-/-^ and *Ldlrad4*^-/-^ mice. Dot plots show yield load, maximum load and stiffness. For each variable in **b, c** and **e** the mean (solid center lines), ±1.0 SD (dotted lines) and ±2.0 SD. (grey boxes) for WT mice (n=320) are shown. Individual data-points for each parameter in *Auts2*^+/^, *Cttnbp2*^-/-^, *Dact3*^-/-^ and *Ldlrad4*^-/-^ lines are shown as green, pink, blue and orange dots, respectively (n=6 animals per genotype). Genes with outlier phenotypes are identified with an asterisk (*) and coloured according to the individual mouse line.

In addition to annotated genes, mice with targeted deletion of the novel, non-annotated genes *Obcdi008175* and *Obcdi042809* also had abnormal skeletal phenotypes, whereas, *Obcdi007392* deficient mice had normal bones (**Supplementary Fig. 11**). Male *Obcdi008175*^-/-^ mice had increased femoral BMC and female *Obcdi008175*^-/-^ mice had decreased femoral length and vertebral strength. By contrast, female *Obcdi042809*^-/-^ mice had decreased vertebral BMC and strength (**Supplementary Fig. 11)**.

These data demonstrate that the *osteocyte transcriptome signature* identifies genes not previously known to affect bone, including novel genes, that play important structural and functional roles in the skeleton.

### *Osteocyte Transcriptome Signature* Genes are Associated with Rare Skeletal Disorders

Given their functional role in mice, we next hypothesised that the *osteocyte transcriptome signature* would be enriched for genes that cause rare monogenic skeletal disorders in humans. Three-hundred and ninety two of 432 skeletal dysplasia-causing gene-orthologs^21^ (∼91%) were actively expressed in osteocytes supporting a key role for osteocytes in skeletal disease. Mutations in 90 genes present in the *osteocyte transcriptome signature* cause 168 of the 612 skeletal disorders annotated in the nosology of genetic skeletal disorders^21^ (3-FE, p=2.4×10^-22^, **Fig. 6a, Supplementary Table 9**). Nevertheless, *osteocyte transcriptome signature* genes were not uniformly involved in all of the skeletal disease groups (**Fig. 6b**). For example, the ‘*osteogenesis imperfecta and decreased bone density group*’ was one of the groups most enriched with signature genes (18/33 casual genes in the signature, p=7×10^-13^). Indeed, all 19 genes known to cause osteogenesis imperfecta (OI)^57–59^ were actively expressed in osteocytes and 14 were present in the *osteocyte transcriptome signature* (**Fig. 6c**). Analysis of the temporal expression pattern of the 19 OI-genes in the IDG-SW3 *in vitro* cell differentiation dataset showed that 16 (84%) were expressed in late osteocytes relative to earlier stages in differentiation, including 13 *osteocyte transcriptome signature* genes (**Supplementary Fig. 12a**). OI-genes were also more highly expressed in osteocytes isolated by laser capture microdissection compared to bone-lining cells and osteoblasts, as well as other tissues (**Supplementary Fig. 12b,c**). Furthermore, 13 of the 14 OI-genes, present in the *osteocyte transcriptome signature,* were up-regulated in osteocytes during skeletal growth but down-regulated at skeletal maturity, suggesting that these genes may have a role in osteocytes during bone development (**Supplementary Fig. 12d**). Together these data demonstrate that the *osteocyte transcriptome signature* is highly enriched for genes that cause rare monogenic skeletal disorders. Thus, interrogation of the *osteocyte transcriptome signature* may inform candidate gene prioritization in the study of families with rare bone disease in which the genetic basis is unknown.

**Fig. 6:**
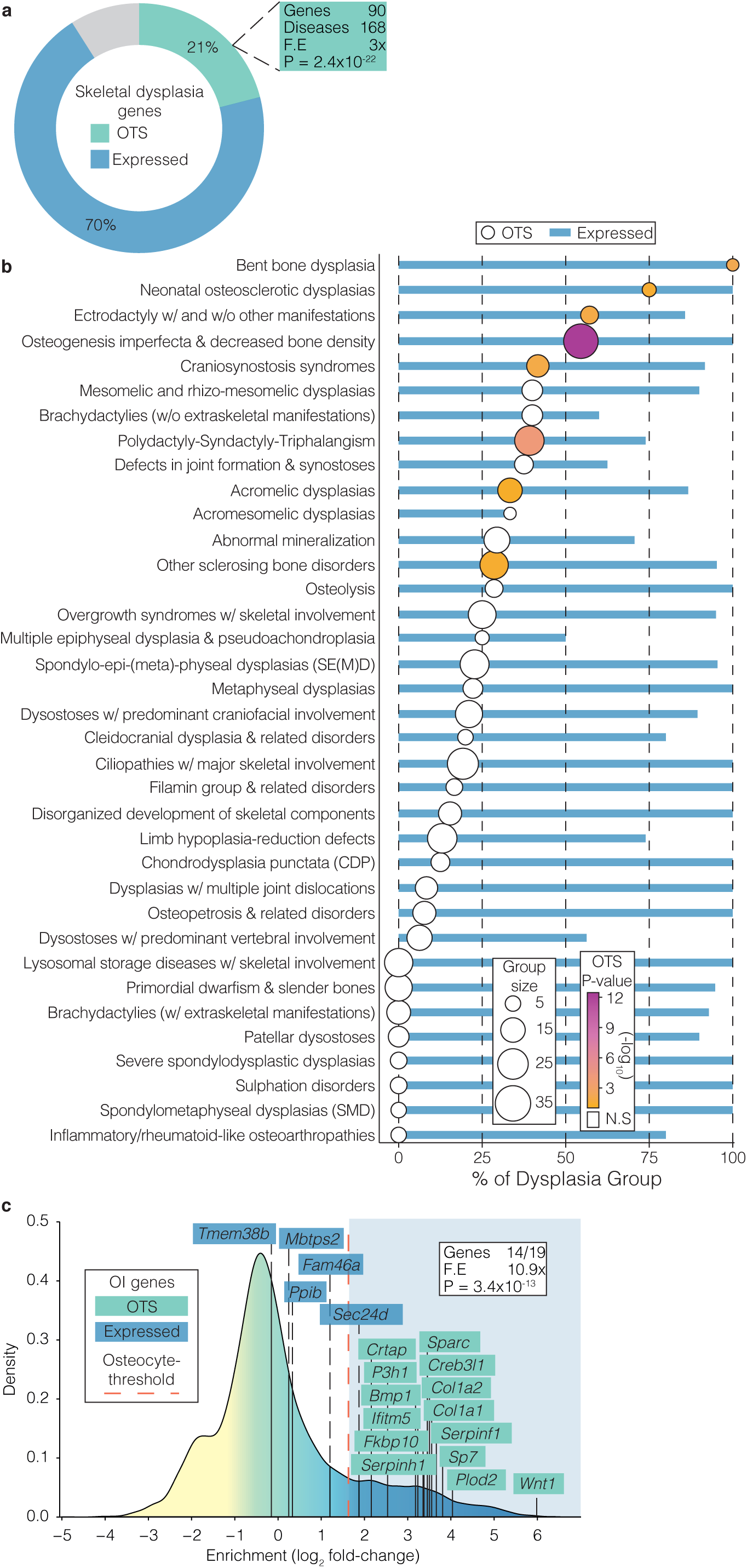
The *osteocyte transcriptome signature* is enriched with genes associated with rare genetic skeletal disorders. **a,** Proportion of genes (%) known to cause genetic skeletal disorders^21^ that have human orthologs either actively expressed in osteocytes (blue), present in the *osteocyte transcriptome signature* (green), or not active in osteocytes (grey). P-value and fold-enrichment (FE) of *osteocyte transcriptome signature* genes among disease-causing genes are calculated under the hypergeometric distribution. **b,** Proportion of genes (%) in individual skeletal disease groups that have orthologs actively expressed in osteocytes (blue bars) or present in the *osteocyte transcriptome signature* (circles). Group size = number of genes in each disease group. Groups with genes significantly enriched in the *osteocyte transcriptome signature* are colored according p-value. Only groups with ≥3 genes are represented. **c,** Osteocyte-enrichment of genes associated with osteogenesis imperfecta (OI). Genes are colored according to their identification as being actively expressed in osteocytes (blue) or present in the *osteocyte transcriptome signature* (green). Bonferroni corrected p-value and fold-enrichment (FE) of *osteocyte transcriptome signature* genes among OI-causing genes are calculated under the hypergeometric distribution.

### The *Osteocyte Transcriptome Signature* is Enriched for Human Orthologs Associated with Skeletal Disease Susceptibility

Finally, we hypothesized that human orthologs of *osteocyte transcriptome signature* genes, identified in mice, are also enriched for genes associated with susceptibility to common skeletal diseases, such as osteoporosis and OA. To test this, the relationship between genetic variations surrounding human orthologs of osteocyte transcriptome signature genes and quantitative ultrasound-derived heel bone mineral density (eBMD) in a sample of 362,924 individuals from the UK Biobank (UKBB) cohort^25^ was examined using two methods: stratified linkage disequilibrium score regression (LDSC-SEG)^60, 61^ and competitive gene set enrichment analysis^62^. LDSC-SEG analysis provided robust evidence of enrichment, demonstrating that genomic regions surrounding human *osteocyte transcriptome signature* gene orthologs contribute disproportionately to the SNP-heritability of eBMD. The *osteocyte transcriptome signature* gene annotation (±20kb) spanned 5.1% of the genome (i.e. 65,332 / 1,142,435 SNPs) and explained ∼12.5% (SE=0.012, P=1.6×10^-7^) of the total estimated SNP-heritability of eBMD (estimated previously^25^ at h^2^_SNP_=0.36, SE=0.017). This corresponded to a 2.2-fold (SE=0.21, P=1.6×10^-7^) enrichment for per-SNP heritability of eBMD.

Competitive gene-set analysis also detected strong enrichment and showed that, on average, human orthologs of *osteocyte transcriptome signature* genes exhibit stronger associations with eBMD than non-*osteocyte transcriptome signature* genes (P=1.8×10^-13^). Enrichment was largely attributable to significant gene-level associations (P<2.6×10^-6^) of 259/992 *osteocyte transcriptome signature* orthologs (26%) with eBMD (**Fig. 7a, Supplementary Table 10**). Mutations in 36 of these genes, with gene-level associations cause monogenic skeletal dysplasia in humans (5.1-fold enrichment, P=5.1×10^-16^) (**Supplementary Table 10**). We observed robust associations between *SOST*, *DMP1* and *MEPE* (P_JOINT_ < 2×10^-129^ P_JOINT_ = 2×10^-27^ and P_JOINT_ = 2×10^-86^, respectively), genes enriched for expression in osteocytes. Given that these genes were reported as the closest gene to a lead variant in previous GWAS of eBMD^25^, we next investigated whether osteocyte transcriptome signature genes occurred nearest to lead variants more often than expected by chance. 110 *osteocyte transcriptome signature* orthologs were the closest genes to conditionally independent lead eBMD variants (2.8-fold enrichment, P=2.4×10^-23^) (**Fig. 7a, Supplementary Table 10**). These included 9 of the top 26 genes most enriched for expression in the *osteocyte transcriptome signature* (*SOST*, *MEPE*, *NGEF*, *WNT1*, *ACKR3*, *TGFB2*, *SEMA3E*, *IRX5*, *DMP1)*. Finally, we identified *AUTS2* (P_JOINT_ =6×10^-9^) as an *osteocyte transcriptome signature* gene that has not previously been implicated by GWAS (i.e. not within 1Mb of a lead eBMD variant), yet reached gene-wide significance in gene-level analysis and caused structural and/or functional skeletal abnormalities when deleted in mice (**Fig. 5a-e**). *LDLRAD4* and *CTTNBP2* (P_JOINT_ =2×10^-20^ and P_JOINT_ =2×10^-87^ respectively), which occur within 1Mb of previously identified eBMD loci, were also shown to result in an abnormal skeletal phenotype when deleted in mice. Together, this demonstrates that variants in osteocyte transcriptome signature gene-orthologs account for a significant proportion of the genetic variance that regulates eBMD and can help identify genes that affect skeletal structure and function.

**Fig. 7:**
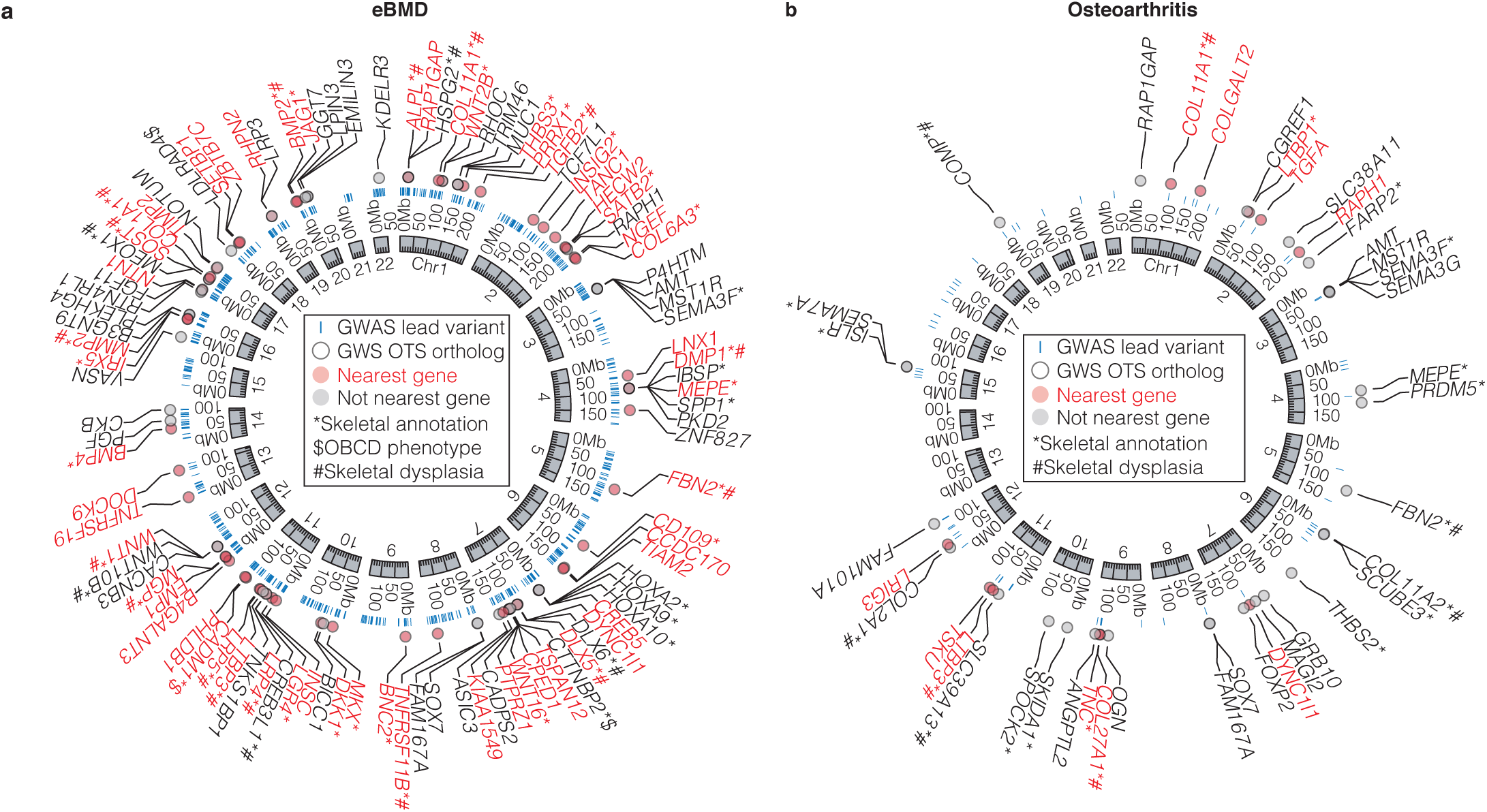
*Osteocyte transcriptome signature* genes are associated with eBMD and osteoarthritis in humans. **a,** The top 100 *osteocyte transcriptome signature* orthologs with significant gene-wise associations (GWS OTS ortholog) with eBMD^25^ (ranked by P_JOINT_< 2×10^-6^). The genome coordinates of each human ortholog are shown. **b,** The 40 *osteocyte transcriptome signature* orthologs with significant gene-wise associations (GWS OTS ortholog) with OA (any subcategory)^64^ (P_JOINT_< 2×10^-6^). Blue bars show the genome coordinates of significant, conditionally independent GWAS variants associated with eBMD (**a**) or OA (**b**) (GWAS lead variants). Genes with significant gene-wise associations and that are located nearest to these loci are shown in red, genes that are not nearest to these loci are shown in black. Genes annotated with a role in the skeleton (either GO biological processes database or skeletal phenotype in the MGI database) are denoted with *, genes in the OBCD database are denoted with $, genes that cause skeletal genetic disorder when mutated in humans denoted with #.

Since changes in subchondral bone structure and mineralization are pathognomonic in the development of OA^63^ we also sought to determine whether variants in *osteocyte transcriptome signature* orthologs were associated with OA in humans. A similar analytical approach was applied to 77,052 individuals with OA and 378,169 control subjects drawn from the UKBB and arcOGEN resources^64^. LDSC-SEG analysis^60, 61^ showed that loci associated with human *osteocyte transcriptome signature* gene orthologs contribute to the heritability of OA at any site (P=1.6×10^-7^), at the knee (P=1.9×10^-6^), hip (P=4.5×10^-3^) and hip and/or knee (P=3.2×10^-6^). Variants associated with *osteocyte transcriptome signature* orthologs explained ∼11.6% (SE=0.011) of the total estimated SNP-heritability of OA at any site. Competitive gene-set analysis^62^ also showed that human orthologs of *osteocyte transcriptome signature* genes had stronger associations with OA than non-osteocyte genes (OA at any site (P=9.1.6×10^-3^), at the knee (P=7.2×10^-3^), hip (P=3.0×10^-3^) and hip and/or knee (P=6.6×10^-2^). Enrichment was largely attributable to significant gene-level associations (P<2.6×10^-6^) of a small number of *osteocyte transcriptome signature* orthologs (40/992, ∼4%) with OA (**Fig. 7b, Supplementary Table 11**). Mutations in 8 of these genes cause monogenic skeletal disorders in humans (7.4-fold enrichment, P=9.5×10^-6^) (**Supplementary Table 11**). 13/40 *osteocyte transcriptome signature* orthologs were the closest genes to conditionally independent lead OA variants (4-fold enrichment, P=1.2×10^-5^), indicating osteocyte transcriptome signature genes occur nearest to lead variants more often than expected by chance (**Fig. 7a, Supplementary Table 11**). Whilst a number of these genes are also expressed in chondrocytes, others such as *MEPE, TSKU, SEMA3F, SEMA3G* and *SEMA7A* (P_JOINT_ <2×10^-6^ in all cases, **Fig. 7b**) are not^65^, indicating their contribution to OA may at least in part be due to their role in osteocytes. Together, these data show that *osteocyte transcriptome signature* genes are not only associated with osteoporosis susceptibility, but may also identify genes associated with OA susceptibility.

## Discussion

Osteocytes are critical cellular regulators of the skeleton. To understand the molecular pathways that control the osteocyte network we generated a map of the osteocyte-enriched transcriptome using data derived from long bones at differing anatomical locations, various ages and both sexes and defined an *osteocyte transcriptome signature* that represents a profile of genes enriched for expression in osteocytes. The majority of these genes have not previously been shown to have a role in bone, including genes that resulted in abnormal structural and functional skeletal phenotypes when deleted in mice. This included novel, non-coding genes, that were restricted in expression to osteocytes, suggesting an additional, unappreciated, level of control over osteocyte function. Integrating this map with orthogonal gene expression datasets^13, 15, 16, 28, 50^, functional skeletal phenotyping data^66^, GWAS datasets^25, 64^, and the nosology of genetic skeletal disorders^21^, provided new understanding of the fundamental role of the osteocyte in skeletal health and disease.

First, we showed the osteocyte transcriptome is conserved among bones from different anatomical locations, yet *Homeobox* genes demarcate osteocytes obtained from upper and lower limbs. This indicates that the molecular ‘postcode’ in osteocytes established during embryonic limb patterning persists in the adult osteocyte, suggesting that their function requires information on their anatomical location. In support of this, site-specific patterns of *Homeobox* genes in adult skeletal progenitors and cells lining bone have been implicated in bone repair^67, 68^, and the regulation of bone mass^69^. Second, we demonstrate the osteocyte transcriptome is regulated differentially between the sexes but only in adulthood, suggesting that osteocyte function during skeletal growth and development is similar in males and females. The adult sexual dimorphism involves genes that control perilacunar remodeling, a process associated with rapid calcium release from the skeleton during lactation, and glucocorticoid-induced bone loss^15, 37, 70^. This expands our understanding of the genes that control this pivotal process. Third, the *osteocyte transcriptome signature* is enriched with genes that control axonal guidance and neuronal network formation, which are up-regulated early in osteocyte differentiation. The remarkable physical and molecular similarity between the osteocyte network and neuronal networks indicates that osteocytes have repurposed neuronal molecular control pathways to facilitate osteocyte network formation and function. Leveraging knowledge from neuronal networks is likely to accelerate understanding of how the osteocyte network forms and functions at a molecular level.

An important challenge facing genome-wide association studies of skeletal diseases is how to map genome-wide significant variants reliably to their causal genes. Integrative analysis of the *osteocyte transcriptome signature* with human genetic association studies of osteoporosis identified new candidate causal genes that may be associated with skeletal disease susceptibility. These included *AUTS2*, *CTTNBP2* and *LDLRAD4,* whose deletion in mice results in abnormal skeletal phenotypes. AUTS2 and CTTNBP2 are regulators of neuronal cell migration and branching^52, 56^, further highlighting the importance of neuron-like pathways in osteocyte network function and in the control of the skeleton. This strategy also identified new candidate loci associated with OA, suggesting that expression of these genes in osteocytes may contribute to the remodeling of subchondral bone, which is critical in the pathogenesis of OA^71^. In addition to complex diseases, analysis of the *osteocyte transcriptome signature* also revealed that genes known to cause monogenic skeletal disorders are enriched in osteocytes. Enrichment was most striking among the ‘*osteogenesis imperfecta and decreased bone density group*’, which is a group characterized by bone fragility. Indeed, the majority of the genes that cause OI are found in the *osteocyte transcriptome signature.* These genes are involved in matrix synthesis and transcript levels were as high or in some cases higher than osteoblasts or bone lining cells in the orthogonal dataset analysis. Whilst these genes are actively transcribed in osteocytes they may not translated until required, for example during perilacunar remodeling. Moreover, these results suggest that, in addition to the canonical model of OI as a disease of osteoblasts, the pathogenesis of OI may also crucially involve the osteocyte network. This conclusion is supported by studies showing the osteocyte network is dysregulated in patients with OI^72, 73^. Together this illustrates that linking knowledge of the *osteocyte transcriptome signature* and functional phenotyping in mice, with GWAS and/or the nosology of genetic skeletal disorders, identifies genes associated with skeletal disease in humans and helps prioritize genes for further analysis.

This study has limitations. The bone samples investigated are long bones and we have not included spine or calvaria. Samples include both cortical and cancellous bone, although cancellous bone represented only 1-2% of the total bone tissue sampled, suggesting the transcriptome data mainly represents genes transcribed in cortical osteocytes. Furthermore, we cannot exclude the possibility that contaminating cells may contribute to the osteocyte-enriched transcriptome, although careful tissue processing and validation using orthogonal datasets strongly suggests the dataset is restricted to genes that are enriched in osteocytes and highly unlikely to include genes expressed only in minor populations of non-osteocyte cells. Lastly, although this study has analysed the trancriptome of osteocytes it has not determined protein expression, nevertheless, skeletal phenotyping of mice with deletions of identified genes have significant skeletal phenotypes suggesting they are translated and functionally important in the skeleton.

Diseases affecting skeletal development, maintenance and repair result in a considerable health burden^74^ and provide the imperative to understand the pivotal role of osteocytes in skeletal physiology and pathophysiology. The osteocyte-enriched transcriptome map and *osteocyte transcriptome signature,* reported here, provide major new insights into the genes and molecular pathways that regulate osteocyte differentiation, osteocyte network formation and mature osteocyte function and are highly enriched for genes implicated in rare and common polygenetic skeletal disease. Thus, defining the *osteocyte transcriptome signature* represents a critical step forward in understanding the fundamental processes underlying skeletal physiology and the cellular and molecular etiology of human skeletal disease.

## Methods

### Transcriptome sequencing mouse cohorts

Transcriptome sequencing and morphological analyses were performed on wild-type, immune-competent, C57BL6/NTac mice. The Garvan/St Vincent’s Animal Ethics Committee approved all animal experiments (Protocol ID 16/01 and 12/44). Mice were maintained in a specific pathogen-free facility and group-housed (2-5 animals per cage) with continuous access to food and water. None of the mice had noticeable health or immune status abnormalities, and were not subject to prior procedures. Three experimental cohorts were used:

#### Bone comparison cohort

Bone samples were collected from left and right tibiae, femora and humeri of eight 16-week-old male C57BL6/NTac mice as detailed in the ‘Sample collection and *in-situ* osteocyte isolation’ section below (n=16 per bone type, 48 samples total). From each mouse, all samples were collected and processed within 20 minutes of sacrifice. Histology and µCT analysis were performed on all samples collected from the right limbs as detailed in the ‘*Morphological analysis of bone samples section’* below (n=8 per bone type, 24 samples total). Bones from the left limbs were processed to obtain *in-situ* isolated osteocytes as detailed in the ‘*Sample collection and in-situ osteocyte isolation’* section below. Transcriptome sequencing was performed on all samples collected from the left limbs as detailed in the ‘*RNA extraction, transcriptome library preparation and RNA-sequencing*’ section below (n=8 per bone type, 24 samples total). Samples were sequenced to an average depth of ∼30 million reads per sample.

#### Skeletal maturation cohort

Left and right humeri were collected from 4, 10, 16 and 26-week-old female and male C57BL6/NTac mice (n=5 per sample type, 80 samples in total). Breeding was stratified so samples from each age could be collected within a single 36-hour time period. Samples were collected in groups of 8 mice (one from each time point in each sex) to avoid confounding batch effects. All samples were collected within 15min of sacrifice. Intact bones from the right limb were used for morphological analysis by DXA as detailed in the ‘*Morphological analysis of bone samples section*’ below. Transcriptome sequencing was performed on all samples collected from the left limb as detailed in the ‘RNA extraction, transcriptome library preparation and RNA-sequencing’ section below (n=5 per bone type, 40 samples total). Samples were sequenced to an average depth of ∼25 million reads per sample.

#### Osteocyte enrichment cohort

Left and right humeri were collected from five 10-week-old male C57BL6/NTac mice (n=5 per sample type, 10 samples total). All samples were collected and processed within 20min of sacrifice. Bones from the left limb were processed to obtain *in-situ* isolated osteocytes as detailed in the ‘*Sample collection and in-situ osteocyte isolation*’ section below. Bones from the right limb were processed in an identical manner but not flushed with PBS or centrifuged so as to retain the bone marrow. Transcriptome sequencing was performed on all samples as detailed in the ‘*RNA extraction, transcriptome library preparation and RNA-sequencing*’ section below (n=5 per sample type, 10 samples total). Samples were sequenced to an average depth of ∼20 million reads per sample.

### Sample collection and *in-situ* osteocyte isolation

Mice were sacrificed by CO_2_ asphyxiation and cervical dislocation. To isolate osteocytes within the bone samples, soft tissue including muscle, ligaments, tendon and periosteum were removed. Diaphyseal bone from the tibia was isolated cutting at the fibula junction and then 1mm distal to the proximal and distal growth plates, and from the femur by cutting the bone immediately proximal to the third trochanter and then 1mm proximal to the distal growth plate. The humeri were cut immediately proximal to the deltoid tuberosity and then 1mm proximal to the epicondyles before completely removing the deltoid tuberosity along the bone shaft. Bone marrow from each bone was removed by first flushing with PBS until visibly clean and then centrifugation at 14,000rcf for 15 seconds. Bones were cut into pieces and snap frozen in liquid N_2_ for storage.

### Morphological analysis of bone samples

#### Dual energy X-ray absorptiometry (DXA)

To examine changes in bone structure in the *Skeletal maturation cohort*, bones were scanned by DXA. Whole femoral length, bone mineral density (BMD) and bone mineral content (BMC) were measured in excised left femora using a Lunar Piximus II dual X-ray absorptiometer (DXA) (GE Medical Systems). Femora were scanned with tibiae attached and the knee joint in flexion to ninety degrees to ensure consistent placement and scan of the sagittal profile.

#### Micro-CT (µCT) visualization of bone samples

bones were scanned using a Skyscan Model 1172 microCT scanner (Bruker) at 50 kV, 200 mA with a 0.5-mm aluminium filter at a pixel size of 4.3 µm. Images were captured every 0.4 degrees through 180 degrees, reconstructed and cortical and cancellous bone regions were defined and bone volume analyzed using NRecon and CTAn software (Bruker, http://bruker-microct.com/products/downloads.htm). Three-dimensional models were created using the Drishti-2 tool^75^ (https://github.com/nci/drishti).

#### Histology

Histological analysis was performed on bone samples from the *Bone Comparison Cohort* to establish the efficacy of the osteocyte isolation methodology. Samples were decalcified in 0.5M EDTA at 37°C for 24 hours and embedded in paraffin. 3μm sections (parasagittal plane) were cut on a RM2265 microtome (Leica), mounted on superfrost plus (Thermo Fisher Scientific, 4951PLUS4) and stained with Mayer’s hematoxylin and eosin (Sigma, MHS1). Images of each section were captured using 10x and 20x objectives with an Aperio Scanscope slide scanner (Leica) and processed by Aperio Imagescope (Leica, https://www.leicabiosystems.com/digital-pathology/manage/aperio-imagescope) and Fiji/ImageJ software^76^ (https://fiji.sc).

#### Tartrate resistant acid phosphatase staining

EDTA-decalcified bone sections (3μ deparaffinized, hydrated and incubated in 1M Tris-HCl pH9.4 buffer at 37°C for 30 minutes. Sections were then stained for Tartrate Resistant Acid Phosphatase (TRAP) by incubation in 1M sodium acetate (pH 5.2), Naphtol-ASBI-phosphate, and sodium tartrate for 10 minutes at 37°C. Sections were then rinsed in distilled water and counterstained with hematoxylin for 15 seconds.

#### Histomorphometry

Histomorphometric assessment of cell-types in cortical bone and on the endocortical surfaces were measured using Osteomeasure software (version 3.2.1.8, Osteometrics Inc). Cell numbers were measured over a 5mm length of both anterior and posterior endocortical surfaces in each bone sample. Measurements began 0.25mm from the first field of view below the proximal end of each sample. Bone cell-types quantified included osteoblasts/bone-lining cells, TRAP positive osteoclasts and osteocytes. Cells that could not be clearly defined as osteocytes, osteoblasts or bone lining cells were defined as ‘other cell-types’.

### RNA extraction, transcriptome library preparation and RNA-sequencing

TRIreagent (Sigma-Aldrich, T9424) was added directly to frozen bone samples and homogenised using a Polytron hand-held homogeniser (PT1200E, Kinematica). RNA was isolated according to the manufacturers protocol and cleaned with an additional ethanol-precipitation step. RNA yield was determined using a Nanodrop (Thermo Fisher Scientific, 2000) and RNA integrity determined using the Bioanalyser RNA 6000 Nano Kit (Agilent Technologies, 5067-1511). Total-RNA (250ng) was depleted of ribosomal RNA using RNaseH (Epicentre) and ribosomal RNA targeting oligonucleotides based on a protocol by Adiconis et al., 2013^77^. Briefly, total-RNA, spiked with External RNA Controls Consortium (ERCC) internal controls (Thermo Fisher Scientific, 4456740), was incubated with ribosomal-RNA targeting oligos (sequences reported in ref^77^) and RNaseH to degrade the rRNAs before the oligos were removed with DNase treatment (Thermo Fisher Scientific, AM2238). RNA was re-purified using magnetic beads according to the manufacturers protocol (Beckman Coulter Genomics, A63987). Total-RNA stranded transcriptome libraries were prepared using the TruSeq Stranded Total RNA LT Sample Prep Kit starting from the fragmentation step in the manufacturers protocol (Illumina, RS-122-2201). Paired-end sequencing (2×125-bp) was carried out on a HiSeq 2500 instrument (Illumina) at the Kinghorn Center for Clinical Genomics, Garvan Institute, Sydney, Australia.

### *De novo* transcriptome assembly and filtering

Sequencing read data was pooled for each bone type in the *Bone Comparison Cohort* (described above) and *de novo* transcriptome assembly performed using two different assembly strategies: *ab initio*, using Trinity^78^, and genome-guided, using Stringtie^79^. Only multi-exon (≥2 exons) transcripts assembled by both methods were retained before transcripts assembled in each of the three bones were pooled using Cuffcompare^80^ to generate a non-redundant union set of assembled transcripts. Assembled transcripts with splice patterns matching those in RefSeq^81^ or GENCODE-M5^82^ transcriptome annotations were removed to identify novel transcripts. Remaining transcripts were then filtered based on exon length. Briefly, the mean log_2_-exon-length ±2 standard deviations of GENCODE-M5 annotated exons was calculated. Assembled transcripts containing one or more exons outside of this range were removed. The protein-coding potential of the remaining transcripts was assessed using CPAT^83^ (http://lilab.research.bcm.edu/cpat). To annotate structures arising from novel loci in the genome, those overlapping known annotated transcripts located on the opposite strand were given a “novel_antisense” biotype and given gene IDs beginning with “Obcda”, while transcripts located between known genes were given the “novel_intergenic” biotype and assigned gene IDs beginning with “Obcdi”. Novel transcripts for both known and novel genes possess transcript IDs begin with “TRINITY”. These novel, multi-exon transcripts were then concatenated to the GENCODE-M5 annotation prior to read alignment. Subsequent to this analysis, new gene structures have been predicted. Some contain splice junctions that overlap novel transcripts reported here. In this case, the GENCODE-M13 gene name is contained in brackets beside the unique assigned gene ID.

### Defining the genes actively expressed in osteocytes

Transcriptome data were trimmed of low-quality reads and adaptor sequences using *Trim Galore!* (https://github.com/FelixKrueger/TrimGalore) and aligned to the GRCm38.p3 mouse genome, guided by the GENCODE-M5 transcriptome annotation plus the novel assembled transcripts described above, using STAR^84^ and quantified using RSEM^85^. A threshold of gene activity was calculated based on the bimodal distribution of normalised gene expression as described in Hart et al^27^. Briefly, FPKM values were log_2_-normalised (nFPKM), omitting genes with FPKM = 0. The kernal density estimate (KDE) of these values was then calculated (using Scotts rule of thumb for bandwidth) and the maximum KDE value determined. A Gaussian distribution was then fitted, with the mean (µ) at the KDE maximum and the standard deviation (SD) based on normalised expression values greater than µ. The nFPKM values were then transformed to zFPKM using zFPKM = (nFPKM - µ)/SD. Active expression in a sample was defined as those with > -2.6 zFPKM, the conservative range suggested in the original publication^27^. This generated sample-specific thresholds for gene activity which are listed in **Supplementary Table 1**. Genes were considered ‘actively expressed’ in a given bone sample type if they were above the sample specific gene activity threshold in all biological replicates. The numbers of active genes in each bone type were classed according to the gene biotype defined in the GENCODE-M5 transcriptome annotation.

### Orthogonal validation of gene expression in osteocytes

To validate the active expression of genes in the osteocyte network and to ensure the genes identified were not derived from minor populations of non-osteocytic contaminating cells we used 3 publicly available datasets;

#### Osteocytic cell-line

This transcriptome sequencing dataset profiled data from the IDG-SW3 mouse osteocyte cell line, an *in vitro* model of osteoblast-like cell to osteocyte differentiation^50^ (ArrayExpress accession E-GEOD-54783). These data were originally published as part of a temporal study of osteocyte differentiation. Data from days 3, 14 and 35 represent osteoblast, early-osteocyte and mature osteocyte stages, respectively^86^. Raw data were aligned and quantified using the GENCODE-M5 transcriptome annotation plus the novel assembled transcripts. Genes were considered to be expressed in this dataset if they had a read count >=10 in each replicate (n=3) in every replicate of early or mature osteocytes.

#### Laser-capture micro-dissected osteocytes

This microarray dataset profiled gene expression in laser capture micro-dissected osteoblasts, bone-lining cells and osteocytes^87^ (GEO accession GSE71306). These data were originally published as part of an investigation into the bone cell response to Sclerostin-antibody treatment. Only untreated control samples were used in this analysis. As this dataset was generated from rat bone, mouse-orthologs of rat genes were identified using the biomaRt package^88^. Background expression and control probes were filtered from the data and mean signal intensity calculated for duplicate probes corresponding to a single gene using the oligo and affycoretools packages^89, 90^. Genes present in each sample were determined using the Wilcoxon signed rank-based gene expression detection algorithm (MAS5calls function from the Affy package^91^). Genes were considered to be expressed in this dataset if they were present in more than 50% of osteocyte samples (>20/40).

#### Collagenase-digested osteocytes

This microarray dataset profiled gene expression in osteocytes from primary bone tissue, with cells removed from the bone surface by flushing, centrifugation and serial collagenase digestions^15^ (ArrayExpress accession E-GEOD-23496). This dataset was originally published in a study investigating gene expression changes in the osteocyte network in virgin mice, mice during lactation and mice post lactation. Background expression and control probes were filtered from the data and mean signal intensity calculated for duplicate probes corresponding to a single gene using the oligo and affycoretools packages^89, 90^. Genes present in each sample were determined using the Wilcoxon signed rank-based gene expression detection algorithm (MAS5calls function from the Affy package^91^). Genes were considered to be expressed in this dataset if they were detected in all replicates (n=3) of any experimental condition.

### Analysis of the osteocyte transcriptome between skeletal sites

#### Gene activity

Gene activity in the *Bone Comparison Cohort* was defined as per the methods section ‘*Defining the genes actively expressed in osteocytes’*. Genes expressed above the sample-specific activity threshold in 8/8 replicates of either tibiae, femora or humeri were considered active. Active genes were classed according to gene biotype defined in the GENCODE-M5 transcriptome annotation.

#### Correlation

The Pearson correlation between each sample was calculated based on normalised counts of active genes in any bone type and plotted using the ggplot2 package^92^. The mean correlation between samples for each bone type comparison is also reported.

#### Differential expression

Differential gene expression analysis between bones was performed using the limma package^93^ on the voom-normalised^94^ counts of active genes in the *Bone comparison cohort*. The topTreat function identified differentially expressed genes with evidence of a log-fold change (LFC) > 0.5 between bone types with a false discovery rate (FDR) corrected p < 0.05. Genes with expression restricted to specific skeletal sites were active in at least one bone type (above the sample specific activity threshold in 8/8 replicates of a given bone type) and inactive in another bone type (below the sample specific activity threshold in 8/8 biological replicates).

#### PCA

Principal component (PC) clustering of samples was performed on the scaled FPKM of all active genes active in the *Bone comparison cohort*, and then separately with actively expressed Homeobox genes using the prcomp R-function. The significance of separation each sample type was determined by Hotellings t-test^95^ and plotted with ggplot2.

### Comparison with other organs and tissues

#### Gene activity

To compare the osteocyte transcriptome with transcriptomes from other tissues, sequencing read data were obtained from Zhang et al^28^ (ArrayExpress accession E-GEOD-54652). Gene activity was defined as per the methods section ‘*Defining the genes actively expressed in osteocytes’*. This dataset contains 8 replicates of 12 tissues types collected for a single study under controlled conditions. Genes expressed above the sample-specific activity threshold in 8/8 replicates of either the adrenal gland, aorta, brown fat, brainstem, cerebellum, heart, hypothalamus, kidney, liver, lung, muscle or white fat were considered active. This identified the active genes in 12 non-skeletal organs and tissues.

#### PCA

To compare the transcriptome of osteocytes and other tissues, FPKM of all genes active in any tissue type were quantile-normalized and scaled before principal components (PCs) were calculated and used to cluster samples. Fitting of 95% CI ellipses and plotting of samples using the first two PCs was performed with ggplot2.

#### Gene specificity

To examine the specificity of gene expression in osteocytes relative to these other tissues we used the ‘Tau’ specificity index^29^. Briefly, median log_2_FPKM values were calculated for each tissue for all genes actively expressed in the osteocyte transcriptome (pooling samples from the 3 skeletal sites to get a single median value for osteocytes). These values were quantile-normalized and Tau calculated for each gene in each tissue using the tispec R-package (https://rdrr.io/github/roonysgalbi/tispec). Genes were classified as either low-specificity (Tau < 0.15), moderate-specificity (0.15 ≤ Tau ≤ 0.85) or high-specificity (Tau > 0.85). The density distribution of Tau in each tissue was plotted with ggplot2, using default axis scaling to ensure visualization of density distribution in each tissue.

### Defining changes in the osteocyte transcriptome with skeletal maturation

#### Gene activity

Gene activity in the *Skeletal Maturation Cohort* was defined per the methods section ‘*Defining the genes actively expressed in osteocytes’*. Genes expressed above the sample-specific activity threshold in 5/5 replicates of any age in either sex were considered active. Active genes were classed according to gene biotype defined in the GENCODE-M5 transcriptome annotation.

#### PCA

PCA was performed on the *Skeletal Maturation Cohort* between female samples (any age), male samples (any age) and sexes at each age (4, 10, 16 and 26-weeks-old). For each comparison, PCs were calculated based on the scaled-FPKMs of genes active in any sample type, with the first two principal components then used to cluster samples. Significant differences between centroids was determined by Hotellings t-test. Fitting of 50% CI ellipses and plotting was performed using the ggplot2 R-package.

#### Weighted gene co-expression network analysis (WGCNA)

To identify clusters of genes with highly correlated patterns of gene expression during skeletal maturation, WGCNA was performed on the normalised counts of genes expressed in either sex at any age (4, 10, 16 or 26-weeks-old) using the WGCNA package^96^. Briefly, gene-wise ’connectedness’ was calculated using the bi-weight mid-correlation function (bicor) across all 40 samples in the *Skeletal Maturation Cohort*. A soft-thresholding power of 8 was calculated as gene connectedness resembled a scale-free network (the scale-free topology model fit R>0.9). Next, a weighted, signed network adjacency matrix was calculated, raising the gene-wise correlation coefficient to the soft-thresholding power with a 10% outlier threshold (maxPOutliers = 0.1). A topological overlap matrix was constructed based on network adjacency and calculated matrix dissimilarity. Hierarchical clustering was performed on the dissimilarity matrix to group genes based on their connectedness and clusters of highly connected genes identified using the hybrid cutreeDynamic R-function^97^. Clusters with correlated patterns of expression were merged (cut-height=0.25) leaving 7 clusters (denoted by colours as per WGCNA convention) of highly connected genes with distinct patterns of expression. Genes that were not correlated with each other or with genes in other clusters were allocated to an 8th ’Grey’ cluster.

#### Cluster characterization

Clusters with expression patterns significantly associated with variation in age, sex or both age and sex were then identified. Briefly, the pattern of gene expression within each WGCNA cluster were summarised into eigengene values (EV_cluster_), defined as the first PC of cluster gene expression variance. Line plots produced with Prism (GraphPad, https://www.graphpad.com/scientific-software/prism) were used to visualise the mean and SD of EV in male and female mice. Three linear models were fitted to each cluster (EV_cluster_ ∼ Age, EV_cluster_ ∼ Sex, EV_cluster_ ∼ Age + Sex + Age*Sex) using the lm function of the stats R-package. The goodness-of-fit of each model was tested with the Bayesian Information Criterion (BIC), with the lowest value across the three models taken to be the optimum for each cluster. The adjusted-R^2^ (adj-R^2^) calculated by the lm function was used to evaluate the strength of each models’ association with EV, with adj-R^2^ > 0.6 considered a ‘strong’ association. For genes in each cluster, expression in osteocytes was validated in independent, orthogonal datasets, as per method section *Orthogonal validation of gene expression in osteocytes*. Significantly enriched Gene Ontology (GO) biological processes^40^, Kyoto Encyclopedia of Genes and Genomes (KEGG) pathways^98^ and Disease Ontology (DO) terms^99^ in each cluster were identified using the ClusterProfiler and DOSE packages^100, 101^ (Bonferroni-corrected p<0.05).

#### Analysis of Magenta cluster genes

Heatmaps of *Magenta Cluster* gene expression and line plots of selected genes (top 20 most strongly correlated with EV_Magenta_) were generated using the mean scaled and normalised gene expression counts, calculated across all ages in both sexes. Heatmaps were produced using the gplots package^102^ and line plots produced with Prism (GraphPad). Semantically similar GO terms were identified within the GO biological processes significantly enriched in the *Magenta Cluster* (Bonferroni-corrected p<0.05) using the ReViGO webtool^103^ (http://revigo.irb.hr/). Briefly, redundant GO terms (similarity > 0.9) were removed and multidimensional scaling (MDS) coordinates calculated based on the SimRel semantic similarity algorithm. MDS coordinates were then used to identify clusters of semantically similar GO terms using the mclust package^104^. The optimum number of clusters (4) was selected among models with unequal variance using the Bayesian Information Criterion (BIC). Bar plots were produced with ggplot2. Expression of top-ranked Magenta genes in osteocytes was assessed using orthogonal datasets described in method section *Orthogonal validation of gene expression in osteocytes.* To determine whether the *Magenta Cluster* identified genes associated with perilacunar-remodeling, their expression was examined during lactation in the *Collagenase-digested osteocytes* microarray dataset^15^ (ArrayExpress accession E-GEOD-23496). Eighty-four of the 95 *Magenta Cluster* genes were represented on the microarray. Competitive gene set testing accounting for inter-gene correlation was performed on each skeletal maturation cluster using the CAMERA function of the limma package^105^. Tukey boxplots of gene expression during lactation were generated for each cluster based on the mean zscores of normalised probe intensity, calculated across all conditions.

### Identification of osteocyte-enriched genes

Genes enriched in osteocytes were identified by comparing gene-counts between the samples of isolated-osteocytes and samples retaining bone marrow, described in the *Osteocyte enrichment cohort*. Gene activity in the *Osteocyte enrichment cohort* was defined as per the methods section ‘*Defining the genes actively expressed in osteocytes*’. Genes expressed above the sample-specific activity threshold in 5/5 replicates of either condition were considered active. Counts for active genes were normalised by library size only and the gene-wise log_2_-fold change (LFC) in normalised read count was calculated between conditions. The density distribution of LFC values was calculated using Scotts rule of thumb for bandwidth. This was plotted to reveal multiple local maxima at different levels of osteocyte enrichment. Genes were grouped within these component populations using a Gaussian Mixture Model (GMM) that was fitted to the LFC density distribution between conditions. The optimum number of components (4) was determined using the BIC among models with unequal variance. K-means clustering (k = 4) was then used to determine initiation parameters for Expectation-Maximization fitting of the 4 component GMM using the mixtools R-package^106^. Gene ontology analysis was performed on the top 1000 genes in each cluster, as ranked by posterior probability, using the ClusterProfiler R-package ^100^. An enrichment threshold was calculated at 2 standard deviations above the mean LFC of the second most enriched GMM component (Component 2) to exclude genes likely belonging to suboptimal components. This identified an empirically determined enrichment threshold = 1.63 LFC. Confidence intervals (95%-CI) of the mean LFC for individual genes was then calculated. Genes with a lower LFC-95%-CI above the enrichment threshold were deemed significantly enriched in osteocytes. Scatter plots, GMM diagrams and density plots were visualized using ggplot2.

### Definition of an osteocyte transcriptome signature

Osteocyte-enriched genes with significantly higher expression in either blood, bone-marrow or skeletal muscle relative to osteocyte enriched bone tissue (p<0.05) were identified using differential gene expression calculations associated with the publicly available data reported by Ayturk et al^13^. Volcano plots were generated using the ggplot2 R-package. The *osteocyte transcriptome signature* was defined as: 1) Genes actively expressed in osteocytes from any of the *Bone comparison cohort*, *Skeletal maturation cohort* or *Osteocyte enrichment cohort* sample types (detailed in section ‘*Defining the genes actively expressed in osteocytes’*) AND 2) Genes enriched for expression in osteocytes relative to bone marrow cells (detailed in section *Identification of osteocyte enriched genes*) AND 3) Genes not significantly enriched for expression in blood, bone marrow or muscle relative to bone. The specificity of osteocyte transcriptome signature gene expression relative to 12 other non-skeletal tissues, was determined using the Tau specificity index for all signature genes calculated as per method section ‘*Comparison with other organs and tissues’*. The histogram of signature gene specificity was generated using ggplot2. To compare expression of *osteocyte transcriptome signature* genes in osteocyte with other bone-cell types we used publicly available microarray dataset which profiled gene expression in laser capture micro-dissected osteoblasts, bone-lining cells and osteocytes^87^ (GEO accession GSE71306). The design and preprocessing of this dataset is described in method section *Orthogonal validation of gene expression in osteocytes – Laser-capture micro-dissected osteocytes*. CAMERA gene-set analysis identified significant differences in expression of signature genes between osteocytes and osteoblasts, and between osteocytes and bone-lining cells. Tukey boxplots were generated based on the mean of scaled, normalised probe intensity values for signature genes in each cell-type using ggplot2.

### Identification of *osteocyte transcriptome signature* genes known to affect the skeleton

*Osteocyte transcriptome signature* genes associated with biological processes important in the skeleton were identified using a curated list of GO biological processes^40^ directly related to the skeleton^107^. Briefly, this list was constructed by filtering GO term descriptions using bone-related keywords. *Osteocyte transcriptome signature* genes associated with any of the 116 manually curated skeletal biological processes were then identified. Similarly, to identify *osteocyte transcriptome signature* genes that cause a significant skeletal phenotype when knocked out in mice, a list of mammalian phenotype (MP) terms related to the skeleton was constructed. MP term definitions and descriptions were filtered using bone-related keywords to identify skeletal MP terms. Screening the Mouse Genome Informatics (MGI) database^41^ with skeletal MP terms identified mouse knockout lines with significant skeletal phenotypes (conditional alleles were excluded). *Osteocyte transcriptome signature* genes associated with a skeletal phenotype when deleted in mice were then identified. Significant over-representation of *osteocyte transcriptome signature* genes in each of these skeletal gene lists was tested under the hypergeometric distribution (p<0.05).

### *Osteocyte transcriptome signature* enrichment and GO semantic similarity clustering

Significantly over-represented GO biological processes and KEGG pathways in the *osteocyte transcriptome signature* were identified using the clusterProfiler R-package^100^ (Bonferroni corrected p<0.05). Clusters of semantically similar GO biological process in the *osteocyte transcriptome signature* were identified as described in the section ‘*Detailed analysis of Magenta Cluster genes*’. The optimum number of clusters (8) was selected among models with unequal variance using the BIC. *Wnt-signaling*, *PTH-signaling* and *Axon-guidance* pathway diagrams coloured by gene expression in osteocytes were constructed using output from the Pathview R-package^108^ and data from the KEGG database^98^.

### *Osteocyte transcriptome signature* gene cluster identification during differentiation

To examine the expression of *osteocyte transcriptome signature* genes during osteocyte differentiation, we analyzed publicly available transcriptome sequencing data from an *in vitro* model of osteoblast-like cell to osteocyte differentiation^50^ (ArrayExpress accession E-GEOD-54783). The design and preprocessing of this dataset is described in method section *Orthogonal validation of gene expression in osteocytes – Osteocytic cell-line*. *Osteocyte transcriptome signature* genes were clustered based on their expression in this dataset, with the optimum number of clusters calculated based on the Calinski-Harabasz (CH) index using the clues R-package^109^ and visualized using the heatmap.2 function of the gplots R-package. Tukey boxplots of cluster expression in each differentiation stage were generated based on the mean zscores of normalised gene expression calculated across all conditions using the ggplot2 R-package. Significant enriched GO biological processes associated with genes in each cluster were identified using the ClusterProfiler R-package (Bonferroni adjusted p<0.05).

### Novel *osteocyte transcriptome signature* gene analysis and deletion in mice

Novel genes were identified based on the *de novo* transcriptome assembly pipeline described in the ‘*De novo transcriptome assembly and filtering’* section above. The gene structures of novel loci and read-data alignment diagrams were visualised using Gvis^110^, using pooled read data from each bone type. Expression in other tissues was determined using the dataset described in section ‘*Comparison with other organs and tissues’*, with bar plots generated with ggplot2.

*Obcdi008175^-/-^*, *Obcdi007392^-/-^* and *Obcdi042809^-/-^* mice were produced by the Mouse Engineering Garvan/ABR (MEGA) Facility (Moss Vale and Sydney, Australia) by CRISPR/Cas9 gene targeting in C57BL/6J mouse embryos using established molecular and animal husbandry techniques^111^. Experiments performed at the MEGA were approved by the Garvan/St Vincent’s Animal Ethics Committee (Protocol ID 18/36). In each case, two single guide RNAs (sgRNAs) were designed to target either side of the genomic DNA encoding the longest predicted transcript and co-injected with polyadenylated Cas9 mRNA into C57BL/6J zygotes. Microinjected embryos were cultured overnight and introduced into pseudo-pregnant foster mothers. Pups were screened by PCR and Sanger sequencing of ear-punch DNA and founder mice identified that carried deletions including the entire gene sequence. The targeted allele was maintained and bred to homozygosity on a C57BL/6J background. The details for each line are as follows:

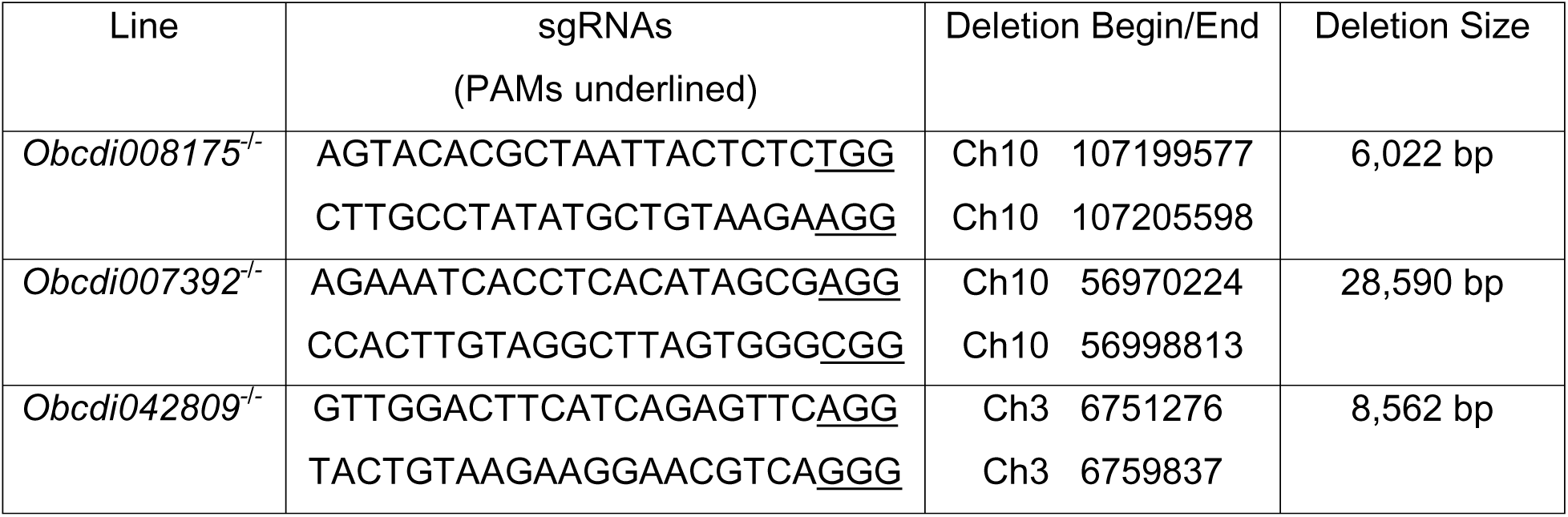

Skeletal phenotyping of these lines was performed as described below (section ‘*Structural and functional skeletal phenotype analysis of mice with deletions of osteocyte transcriptome signature genes*’) except that digital X-ray images were recorded at a 10 m resolution using a 10 μ m resolution using a Faxitron UltraFocus (Faxitron Bioptics LLC,Tucson, Arizona USA) operating in full manual mode at 22kV.

### Structural and functional skeletal phenotype analysis of mice with deletions of *osteocyte transcriptome signature* genes

The Origins of Bone and Cartilage Disease (OBCD) program^66^ is undertaking a validated rapid throughput multiparameter skeletal phenotype screen of mutant mouse lines generated by the Wellcome Trust Sanger Institute as part of the International Mouse Phenotyping Consortium (IMPC)^112^ and International Mouse Knockout Consortium (IMKC)^113^ effort. The OBCD mouse studies were undertaken by Wellcome Trust Sanger Institute Mouse Genetics Project (MGP) as part of the IMPC. This was licensed by the UK Home Office (PPLs 80/2485 and P77453634) in accordance with the 1986 Animals (Scientific Procedures) Act and approved by the Wellcome Sanger Institute’s Animal Welfare and Ethical Review Body.

All mice generated by the MGP were produced (Tm1 alleles^114^, CRISPR alleles^115^) and maintained on a C57BL/6NTac background. Mice were fed either a Breeder’s Chow (Mouse Breeder Diet 5021, 21% kcal as fat, Labdiet, London, UK) or a high fat diet (Western RD, 829100, Special Diet Services, Witham, UK) from 4 weeks of age. Prior to tissue collection, mice underwent a standardized broad primary phenotype screen (https://www.mousephenotype.org/impress/PipelineInfo?id=15)^116^. Detailed OBCD phenotyping methods were then performed as below:

#### Origins of Bone and Cartilage Disease Phenotyping Methods

Left lower limb and tail samples from 16-week-old female wild-type and knockout mice were stored in 70% ethanol at 4°C, anonymized and randomly assigned to batches for rapid-throughput analysis in an unselected fashion (n=2-6 per line). Overall, 19 structural and functional skeletal parameters were determined for femur and vertebrae samples from each mouse studied and compared to reference data (shown as a range with mean and 1 and 2 standard deviations) obtained from 320 16-week-old wild-type C57BL/6NTac female mice and collected in batches from across the study time-course. Coefficients of variation for each skeletal parameter were: femur BMC (2.0%) and length (2.1%); vertebra BMC (2.1%) and length (2.3%); trabecular bone volume/tissue volume (18.5%), trabecular number (7.3%), trabecular thickness (7.9%) and trabecular spacing (8.3%); cortical bone thickness (4.3%), internal diameter (6.0%) and BMD (4.0%); femur yield load (13.2%), maximum load (10.0%), fracture load (29.0%), stiffness (13.7%) and energy dissipated before fracture (26.7%); and vertebra yield load (13.0%), maximum load (10.3%) and stiffness (13.3%).

#### Digital x-ray microradiography

Soft tissue was removed from skeletal samples and digital X-ray images were recorded at a 10μm resolution using a Faxitron MX20 operating at 26kV (Qados, Cross Technologies plc, Sandhurst, Berkshire, UK). Cleaned lower limb and caudal vertebrae 6 and 7 were imaged together with 1mm diameter steel, aluminum and polyester standards. Bone lengths and relative bone mineral content (BMC) were determined as previously described^66^. Briefly, femur length and mean length of caudal vertebrae 6 and 7 were determined using ImageJ and images calibrated using an X-ray image of a digital micrometer. To determine relative BMC, 2368×2340 16-bit DICOM images were converted to 8-bit Tiff images in ImageJ, the grey levels of the polyester and steel standards were defined and the image stretched between the polyester (grey level 0) and steel (grey level 255) standards. Increasing gradations of mineralization density were represented in 16 equal intervals by applying a pseudocolour lookup table to each image. For each sample the median grey level (0-***1.*** 255) of the femur and caudal vertebrae 6 and 7 was calculated.

#### Micro-CT Analysis

Cortical and trabecular parameters were determined by micro-CT using a Scanco μCT50 (Scanco medical, Zurich, Switzerland). Samples were scanned at 70kV, 200μA, with a 0.5mm aluminium filter, 1 second integration time, no averaging, and images captured every 0.36° though 180° rotation. Reconstructions, ROI selection and analyses were performed using Scanco software. Trabecular bone parameters (trabecular bone volume as a percentage of tissue volume BV/TV, trabecular number Tb.N, trabecular thickness Tb.Th, and trabecular spacing Tb.Sp) were calculated from scans at a voxel resolution of 5μm in a 1mm long region of the trabecular compartment beginning 100μm proximal to the distal femoral growth plate. Cortical bone parameters (cortical thickness Ct.Th, internal endosteal diameter, and BMD) were calculated from scans at voxel resolution of 10μm from a 1.5mm long region of mid-shaft cortical bone centred 56% along the length of the femur distal to the femoral head.

#### Micro-CT analysis of osteocyte lacunae

Osteocyte lacuna parameters were determined by micro-CT using a Scanco μCT50 as described above. Osteocyte parameters (lacuna number Lc.N, lacuna volume Lc.V, and lacuna sphericity Lc.Sph) were calculated from scans at a voxel resolution of 1 μm from a 0.25mm long region of mid-shaft cortical bone centred 60% along the length of the tibial from the tibial plateau. The micro-CT segmentation threshold was optimised by analysing lacuna area (Lc.Ar and Lc.N) in longitudinal sections of cortical bone imaged by both back scattered-election scanning-electron microscopy (considered the gold standard) and micro-CT. Micro-CT data was reconstructed using a range of segmentation thresholds and the distribution of Lc.Ar and Lc.N did not differ between BSE-SEM and micro-CT at a segmentation threshold of 350 (equivalent to 851.7 mg HAcm^-3^). Therefore, a segmentation threshold of 350 was subsequently used for the analysis of all samples. DICOM images were processed using Fiji^76^ and lacunae rendered using its Volume Viewer (https://imagej.nih.gov/ij/plugins/volume-viewer.html). Lc.V was determined using BoneJ Particle Analyzer^117^ and L.Sph using Fiji 3D Shape Measure^118^. Following determination of Lc.N and Lc.N/BV for each sample, the maximum equivalent number of lacunae were randomly selected from each sample (2693) and the volume and sphericity distribution for each mutant was compared to WT using Kruskall-Wallis analysis (GraphPad Prism 8). Differences between mutant and WT were considered valid if significant in all 10 permutations performed.

#### Biomechanical Testing

Destructive 3-point bend tests and compression tests were performed on an Instron 5543 materials testing load frame (Instron Limited, High Wycombe, UK). Femur strength and toughness (yield load, maximum load, fracture load, stiffness, % energy dissipated prior to fracture) were derived from destructive three-point bend testing using a 50-N load cell and custom mounts with rounded supports to reduce cutting and shear loads. Bones were positioned horizontally with the anterior surface upwards between two mounting points with a span of 8mm. Load was applied vertically to the mid-shaft with a constant rate of displacement of 0.03mm/second until fracture. The biomechanical properties of caudal vertebrae 6 and 7 (yield load, maximum load and stiffness) were derived from compression testing using a 500-N load cell and two custom anvils. Vertebrae were bonded in vertical alignment to a custom anvil support using cyanoacrylate glue and load was applied vertically at a constant rate of displacement of 0.03 mm/s and a sample rate of 20 Hz until approximately 1mm of displacement had occurred^66^.

### Tissue expression screen

Expression in other non-skeletal organs tissues of *Auts2*, *Dact3*, *Ldlrad4* and *Cttnbp2* and genes with established role in osteocyte biology^38^ was analysed using data described in method section *‘Comparison with other organs and tissues’*. The mean expression in each tissue was first calculated and then the percentage of this mean relative to maximum mean expression across all tissues used to construct heatmaps with the ggplot2 package. Bar plots of normalised expression (FPKM) were constructed with ggplot2.

### Association of *osteocyte transcriptome signature* genes with genetic skeletal disorders

*Osteocyte transcriptome signature* gene orthologs known to cause rare skeletal genetic disorders in humans were identified in the Nosology and Classification of Genetic Skeletal Disorders^21^. Significant over-representation was examined under the hypergeometric distribution, using the parameters:

- Population - the number of mouse genes with human orthologs actively expressed in any of the *Bone comparison cohort*, *Skeletal maturation cohort* or *Osteocyte enrichment cohort*s or any of the 12 tissues described in the *Comparison with other organs and tissues* section (15,368 genes)
- Successes in population - the number of genes known to cause genetic skeletal disorders (either total or within each nosology-defined skeletal disease groups) within the population (432 genes)
- Sample size - the number of *osteocyte transcriptome signature* genes within the population(1,043 genes)
- Successes in sample - the number of *osteocyte transcriptome signature* genes known to cause genetic skeletal disorders(either total or within each individual nosology-defined skeletal disease groups) within the population (90 genes)

Significant enrichment of *osteocyte transcriptome signature* gene orthologs among all genes in the nosology, and within each of the nosology-defined skeletal disease groups was calculated using the Graeber lab online hypergeometric calculator (https://systems.crump.ucla.edu/hypergeometric/index.php). Donut plots, bubble plots and bar charts were constructed using ggplot2. Genes known to cause osteogenesis imperfecta (OI genes) were identified^57^ with the addition of *SEC24D*^59^ and *FAM46A*^58^. OI-gene expression during osteocyte differentiation was assessed using an independent, orthogonal dataset described in method section *Orthogonal validation of gene expression in osteocytes – Osteocytic cell-line*. OI-gene expression in osteocytes relative to other bone cell types was assessed with an independent, orthogonal dataset described in method section *Orthogonal validation of gene expression in osteocytes – Laser-capture micro-dissected osteocytes.* Significant differences in gene expression between cell types were calculated by 2-way ANOVA (adjusted for multiple-comparisons) and visualized using Prism (GraphPad). Expression in other non-skeletal organs tissues was analysed using data described in method section ‘*Comparison with other organs and tissues*’. The mean expression in each tissue was first calculated and then the percentage of this mean relative to maximum mean expression across all tissues used to construct heatmaps with the ggplot2 R-package. OI gene expression during skeletal maturation was analysed using data described in section ‘*Defining changes in the osteocyte transcriptome with skeletal maturation’*. The strength and significance of Pearson-correlation between OI gene expression and age were calculated for each sex using the cor,test function in the stats R-package. Lineplots were produced in Prism (GraphPad).

### Enrichment of osteocyte transcriptome signature for orthologs associated with Osteoporosis and OA

Competitive gene-set analysis, stratified linkage disequilibrium score regression, and analysis of genes nearest to significant loci, were used to investigate whether human osteocyte transcriptome signature orthologs associated with: (i) quantitative ultrasound derived heel bone mineral density (eBMD), a predictor of osteoporosis susceptibility, and (ii) self-reported, or hospital-diagnosed osteoarthritis.

#### Datasets used for analysis

Analyses of eBMD were performed on a sample of 362,924 unrelated white British subjects (54% female, GCTA pairwise relatedness < 0.10) from the UK Biobank Study (UKB)^119^ that had valid quantitative eBMD and high-quality genome-wide HRC and 1000G/UK10K imputed data from the January 2018 release [(i.e. 20,490,436 genetic variants with an information quality score > 0.3, minor allele frequency > 0.05%, minor allele count > 5, genotyping hard call rate > 0.95, and weak evidence of deviation from Hardy-Weinberg equilibrium (P > 1×10^-6^)]. Analyses of individuals with OA were performed on summary results statistics from a recent UKBB and arcoGEN GWAS meta-analysis of four predefined OA subcategories: Osteoarthritis at any site (ALLOA): 77,052 cases & 378,169 controls, hip osteoarthritis (HIPOA): 15,704 cases & 378,169 controls, knee osteoarthritis (KNEEOA): 24,955 cases & 378169 controls, and hip and/or knee (HIPKNEEOA): 39,427 cases & 378,169 controls. Details of how eBMD and OA was defined and criteria for including individuals in each cohort are detailed in the original publications^25, 64^.

***1. Competitive gene set analysis.*** *Overview:* Competitive gene set analysis involved a three-stage process. In the first stage mouse – human orthologues were mapped and the gene universe used for enrichment defined. In the second stage gene-based tests of association were performed using genome-wide genetic data to estimate the strength of association of each human protein-coding gene with eBMD or OA. In the third stage, competitive gene set analysis was used to compare the mean strength of association of human orthologues of osteocyte transcriptome signature genes, to the mean strength of association of non-osteocyte transcriptome signature genes. Evidence of enrichment was obtained through significance testing, in which the null hypothesis of no difference in mean association between osteocyte, and non-osteocyte signature gene sets was tested against a one-sided alternative that stipulated that osteocyte transcriptome signature genes were on average more strongly associated with eBMD or OA than non-osteocyte transcriptome signature genes.

*Stage 1- Orthologue mapping and universe definition:* Mouse-to-human orthologs were identified in the February 2014 Ensembl database archive, in line with the GRCh37 genome used in the GWAS analysis, accessed through the biomaRt R-package^88^. To ensure enrichment was calculated relative to genes that had a fair chance of identification in the osteocyte transcriptome signature, and avoid p-value inflation due to inclusion of genes not able to be assayed by RNA-seq, the gene universe was limited to orthologs of genes actively expressed in any sample type from either of the *Bone comparison*, *Skeletal maturation* or *Osteocyte enrichment* cohorts, or any of the 12 non-skeletal tissues detailed in the *Comparison with other organs and tissues* section. This encompassed 16,015 mouse-human orthologs, including 992 of the *Osteocyte transcriptome signature* genes. All analyses were repeated relative to all human-mouse gene orthologs. In all cases, the results from analyses relative to the gene universe were more conservative than when all orthologs were used, and the more conservative results are presented here.

*Stage 2- Gene-based tests of association:* Gene-based tests of association were conducted in MAGMA (v1.06, https://ctg.cncr.nl/software/magma)^62^ using imputed individual level genotype data for the analyses involving eBMD, and GWAS meta-analysis summary results statistics for OA. Analyses involving eBMD were further adjusted for age, sex, genotyping array, assessment centre and ancestry informative principal components 1 – 20. Gene-based tests of association encompassed a multi-model approach in which the association results from different gene analysis models were combined to produce an aggregate p-value corresponding to the strength of evidence of association between each gene (±20kb) and eBMD or OA. The three association models included: a principal components regression model, a SNP-wise mean χ2 model [*i.e*. test statistic derived as the sum of −log(SNP p-value) for all SNPs that intersect the gene region of interest], and SNP-wise top χ2 model [(test statistic derived as the sum of −log(SNP p-value) for top SNP in the region of interest)]. The aggregate approach was chosen as it yields a more even distribution of statistical power and sensitivity over a wider range of different genetic architectures. Note: Principal components regression model could not be run for OA as the method requires individual level genotyping data that was not available. The enrichment of genes with mutations known to cause human genetic skeletal disorders^21^ was examined under the hypergeometric distribution using the following parameters:

- Population: Orthologs in the above defined gene universe (16,015 genes)
- Total number of successes in the population: Genes in the population known to cause skeletal disorders when mutated in humans (432)
- Sample: Osteocyte transcriptome signature orthologs in the population with significant gene-level associations with either eBMD or OA (eBMD = 259, OA = 40)
- Number of sample successes: The number of Osteocyte transcriptome signature orthologs in the population with significant gene-level association with either eBMD or OA known to cause human skeletal disorders (eBMD = 36, OA = 8)

Significant enrichment was calculated using the Graeber lab online hypergeometric calculator (https://systems.crump.ucla.edu/hypergeometric/index.php).

*Stage 3- Gene set analysis:* Competitive gene set analysis was used to determine whether the set of 992 osteocyte transcriptome signature human gene orthologues was on average more strongly associated with eBMD or OA than non-osteocyte signature genes. The analysis accounted for several confounding factors including: gene size, gene density (i.e. representing the relative level of LD between SNPs in the gene) and the inverse of the mean minor allele count in the gene (i.e. to correct for potential power loss in very low minor allele count SNPs), as well the log value of these three factors.

Human genome coordinates (hg19) were mapped and Circos plots generated for the *osteocyte transcriptome signature* orthologs with significant gene-level associations with eBMD and OA using the ggbio R-package^120^. The top 100 genes associated with eBMD (ranked by p-value) were shown to avoid over-plotting.

***2. Stratified Linkage disequilibrium Score Regression (LDSC-SEG).*** Stratified linkage disequilibrium score regression^60, 61^ was used in conjunction with summary results statistics from recent GWAS of eBMD and OA^25, 64^ to investigate whether genomic regions surrounding *osteocyte transcriptome signature* human gene orthologs contribute disproportionately to the SNP heritability of eBMD and OA. Here, heritability was defined as the proportion of trait variation / disease liability explained by genome-wide imputed genetic markers and is referred to as SNP-heritability. Enrichment is expressed in terms of per-SNP heritability, and estimated as the proportion of SNP heritability explained by genomic regions intersecting the gene set, divided by the proportion of SNPs intersecting the corresponding gene set. Evidence of enrichment is evaluated though significance testing, in which the null hypothesis of no difference in per-SNP heritability, is tested against the one-sided alternative where the per-SNP attributable to the gene-set is greater than the per-SNP heritability attributable to the rest of genes in the genome. Using a similar approach to that described previously^60, 61^, we constructed a genome-annotation for autosomal gene regions ± 20kb of the *osteocyte transcriptome signature* orthologs, a second encompassing all mappable mouse-human genes, and a third annotation corresponding to the gene universe described above. We applied LDSC-SEG^60, 61^ to jointly model the *osteocyte transcriptome signature* annotation, together with 52 functional genomic annotations that include genic regions, enhancer regions and conserved regions (i.e. baseline model v1.1 supplied with the LDSC-SEG software, https://github.com/bulik/ldsc). We limited the analysis to high quality imputed autosomal SNPs (INFO > 0.9), excluded the HLA region from all analyses and used the 1000 Genomes LD reference panel of unrelated European subjects as supplied with the software. Enrichment was quantified by the LDSC-SEG regression coefficient, which corresponds to the magnitude of enrichment in per-SNP eBMD heritability attributable to the *osteocyte transcriptome signature*, conditional on the gene universe and the 52 baseline annotations. Strength of evidence against the null hypothesis of no enrichment attributable to the *osteocyte transcriptome signature* annotation, conditional on other annotations was determined by the LDSC-SEG p-value. Sensitivity analysis was conducted by increasing the window to ±100kb of each gene and re-analysing.

3. ***Nearest-gene enrichment analysis.*** Lastly, we examined whether osteocyte transcriptome signature orthologs were located nearest to genome wide significant (GWS), conditionally independent GWAS loci associated with eBMD and OA more often than would be expected by chance. This was examined under the hypergeometric distribution using the following parameters:

- Population: Orthologs in the above defined gene universe (16,015 genes)
- Total number of successes in the population: Number of osteocyte transcriptome signature orthologs in the population (992)
- Sample: Genes in the population that are the nearest gene to a GWS loci (eBMD = 638, OA = 52)
- Number of sample successes: Number of osteocyte transcriptome signature orthologs in the population that are the nearest gene to a GWS loci (eBMD = 110, OA = 13)

Significant enrichment was calculated using the Graeber lab online hypergeometric calculator (https://systems.crump.ucla.edu/hypergeometric/index.php). Only unique genes were used to calculate enrichment (genes located nearest to multiple GWS loci were only counted once).

### Quantification and statistical analysis

Statistical methodologies and software used for performing these analyses are described in the appropriate sections. Biological replicates were taken from distinct samples. Analyses using Trinity (v2.0.6), Stringtie (v1.0.4), Cuffcompare (v2.2.1), Trimgalore (v0.3.3), STAR (v2.4.1d) and RSEM (v1.2.21) were performed on a computing cluster running the CentOS 6.8 (Rocks 6.2) Linux operating system. CTAn, NRecon and Drishti were run using a Windows 7 OS. CPAT and ReViGo analyses were run using the web interface. Hypergeometric enrichment was tested using the Graeber lab online hypergeometric calculator (https://systems.crump.ucla.edu/hypergeometric/index.php). Gene-based tests of association and competitive gene set analysis were conducted with MAGMA (v1.06, https://ctg.cncr.nl/software/magma). Stratified linkage disequilibrium score regression was conducted with LDSC-SEG software, https://github.com/bulik/ldsc (v1.1). All other statistical analysis was performed in R (>v3.4.0), with key packages cited in the methods text. Error bars reflect mean and standard deviation unless stated otherwise. Multiple hypothesis correction was used wherever significance was evaluated across multiple statistical tests i.e. differential gene expression analysis (Benjamini-Hochberg FDR), GO enrichment, KEGG enrichment, DO enrichment, Nosology group enrichment (Bonferroni correction).

### Data availability

The raw RNA-sequencing data (fastq), read alignment files (BAM) and processed gene expression data files for each cohort (FPKM and counts) are deposited at ArrayExpress (https://www.ebi.ac.uk/arrayexpress) under the following accession numbers: E-MTAB-5532 (*Bone comparison cohort*), E-MTAB-7447 (*Skeletal maturation cohort*) and E-MTAB-5533 (*Osteocyte enrichment cohort*). Publicly available datasets used in this work are available at ArrayExpress or the Gene Expression Omnibus (GEO) accession IDs as indicated in the relevant method sections above.

### Code availability

Analysis scripts are available upon reasonable request to the authors.

## Acknowledgements

This work was supported by a Wellcome Trust Strategic Award (101123/Z/13/A) to G.R.W, P.I.C and J.H.D.B, and by Mrs Janice Gibson and the Ernest Heine Family Foundation. P.I.C is supported by Mrs Janice Gibson and the Ernest Heine Family Foundation. J.H.D.B and G.R.W received a Wellcome Trust Joint Investigator Award (110141/Z/15/Z and 110140/Z/15/Z) and EU H2020 THYRAGE Grant (666869). S.E.Y and A.P.C were funded by the Australian Government Research Training Program Scholarships. J.P.K was funded by a University of Queensland Development Fellowship (UQFEL1718945), a National Health and Medical Research Council (Australia) Investigator grant (GNT1177938) and project grant (GNT1158758). J.T.S was funded by a University International Postgraduate Award Scholarship (UNSW). J.A.M was funded by a Banting Postdoctoral Fellowship. J.M.W.Q was funded by an Australian National Health and Medical Research Council project grant (ID: GNT1057706). M.J was funded by Viapath. J.B.R was supported by the Canadian Institutes of Health Research (CIHR), the Lady Davis Institute of the Jewish General Hospital, the Canadian Foundation for Innovation, the NIH Foundation, Cancer Research UK and the Fonds de Recherche Québec Santé (FRQS). JBR is supported by a FRQS Clinical Research Scholarship. TwinsUK is funded by the Welcome Trust, Medical Research Council, European Union, the National Institute for Health Research (NIHR)-funded BioResource, Clinical Research Facility and Biomedical Research Centre based at Guy’s and St Thomas’ NHS Foundation Trust in partnership with King’s College London. D.J.A and C.J.L were funded by Wellcome Trust Strategic Awards. D.M.E. was funded by an Australian National Health and Medical Research Council Senior Research Fellowship (APP1137714). T.G.P was funded by an Australian National Health and Medical Research Council Senior Research Fellowship (1155678). This research has been conducted using the UK Biobank Resource (accession IDs: 12703). We are grateful for the phenotyping expertise provided by Anne-Tounsia Adoum, Justyna Miszkiewicz, Rebecca Allen, Jayashree Bagchi Chakraborty, Katharine F Curry, Hannah F. Dewhurst, Apostolos Gogakos, Naila Mannan, Hayley J Protheroe, Penny C Sparkes, Valerie Vancollie and Fiona Kussy. We thank Ya Xiao for assistance with animal studies. We thank the Sanger Institute’s Research Support Facility, Mouse Pipelines and Mouse Informatics Group who generated mice and collected materials for this manuscript.

## Author contributions

S.E.Y, J.H.D.B, G.R.W, P.I.C conceived the study. S.E.Y, J.P.K, M.I, A.B-M, F.R, E.D, J.B.R, R.B, T.G.P, J.A.E, D.M.E, E.Z, P.A.B, J.H.D.B, G.R.W, P.I.C designed experiments. S.E.Y, J.G.L, E.J.G, C.M.S, M.D, S.E.F.G, V.D.L, N.C.B, D.K-E, R.C.C, S.M, M.M.M, J.M.W.Q, A.R.M conducted experiments. J.G.L, E.J.G, C.M.S, S.M, M.D, S.E.F.G, V.D.L, N.C.B, D.K-E, R.C.C, D.J.A, C.J.L, R.B generated mouse models and/or functional experiments. S.E.Y, J.P.K, J.G.L, C.M.S, S.M, M.I.D, S.E.F.G, V.D.L, N.C.B, D.K-E, R.C.C, A.P.C, J.T.S, J.A.M, N.B, M.J, K.H, J.H.D.B, G.R.W, P.I.C performed data analysis. S.E.Y was the lead analyst. S.E.Y, J.P.K, J.H.D.B, G.R.W, P.I.C wrote the manuscript. All authors revised and reviewed the paper.

## Supplementary Figure Legends

**Supplementary Fig. 1:**
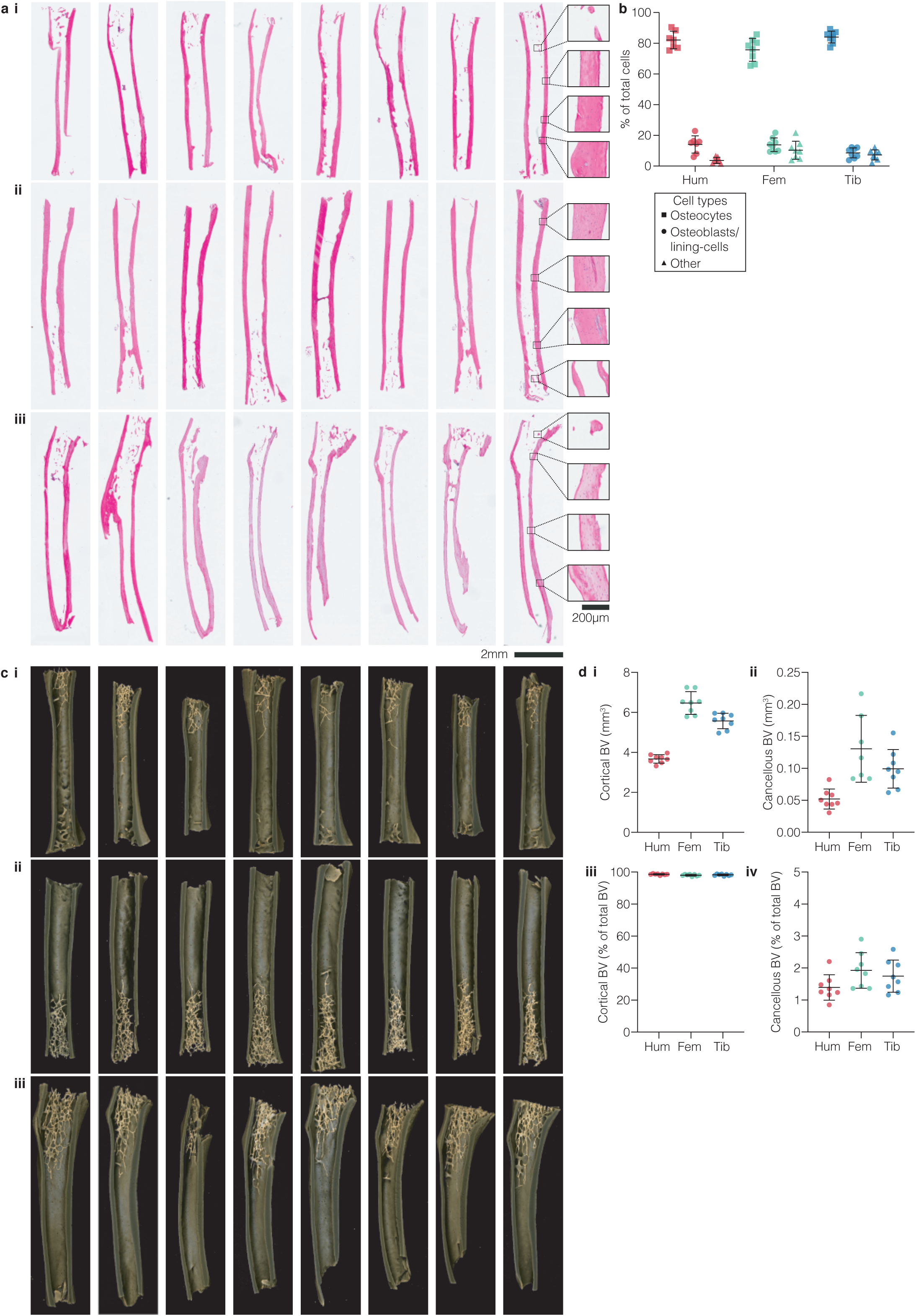
Osteocyte enrichment in processed bone samples. **a,** Histology images of bone samples from the humeri (i), femora (ii) and tibiae (iii). Example high-magnification images show the effective removal of marrow and endosteal cells and enrichment for osteocytes. **b,** Quantitative analysis of bone-cell types present in osteocyte-enriched samples. ‘Other’ denotes all cell types that could not be defined as osteoblasts, lining cells or osteocytes. Empty lacunae were excluded. Individual data points are shown. Each point represents data from a single bone from one mouse. Error bars show mean and SD.**c,** Micro-CT images of osteocyte-enriched bone samples from the humeri **(i)**, femora **(ii)** and tibiae **(iii)** following processing. Samples correspond to those in panel **a** above. Samples were collected from the contralateral limb and processed at the same time as those used for transcriptome sequencing. **d,** The cortical **(i)** and cancellous **(ii)** bone volume (BV) in processed bone samples from each skeletal site, and the relative proportion of total BV (as a percentage) of each bone type **(iii-iv)**.

**Supplementary Fig. 2:**
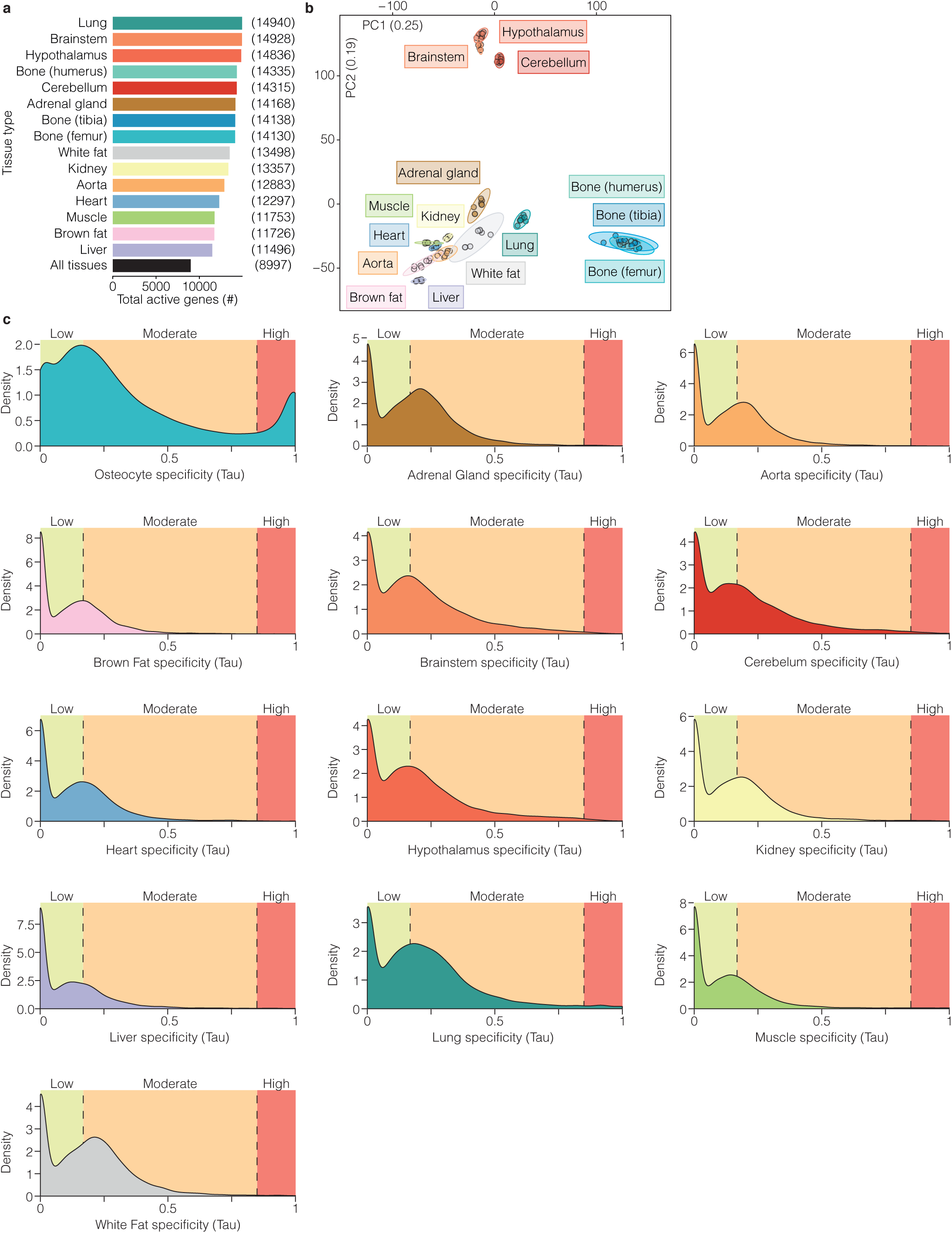
The osteocyte transcriptome is distinct from transcriptomes expressed by other organs and tissues. **a,** The number of genes actively expressed in osteocytes from the humerus, tibia and femur, and 12 other organs and tissues^28^. The black bar indicates the number of genes common to all tissues. **b,** Principal component analysis of active gene expression in osteocytes isolated from the humerus, tibia and femur, and 12 other organs and tissues. Dots represent individual biological replicates and ellipses represent 95% confidence intervals for each sample type. Coloured labels correspond with sample type. The percentage of total variance explained by individual PCs is shown. **c,** The distribution of active gene expression specificity^29^ (Tau) in osteocytes and 12 non-skeletal organs and tissues, calculated for each gene in the osteocyte transcriptome. Genes with Tau < 0.15 have low expression specificity in a given tissue (green), 0.15 ≤ Tau ≤ 0.85 have moderate specificity (orange), while Tau > 0.85 have high expression specificity (red).

**Supplementary Fig. 3:**
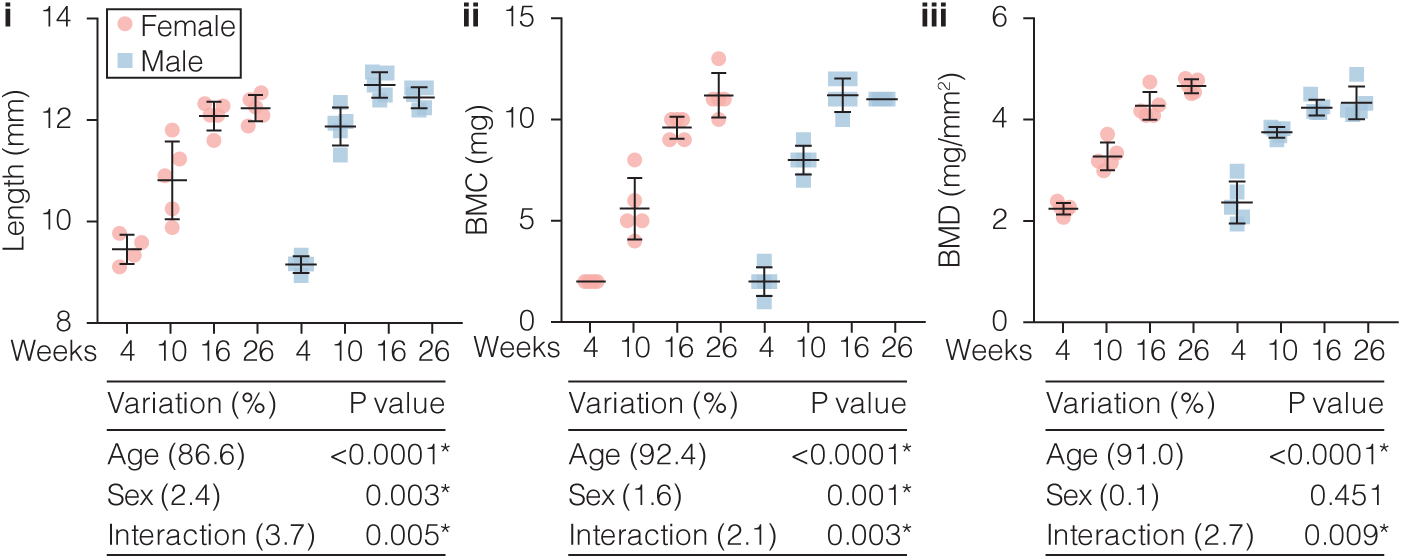
Differences in bone structure during skeletal maturation in both sexes. Bone structural parameters, humeri length **(i)**, bone mineral content (BMC) **(ii)** and bone mineral density (BMD) **(iii)** from female (pink) and male (blue) mice at different ages (weeks). Individual samples are denoted by dots and the mean ±SD are shown. Statistical analysis performed by two-way analysis of variance (ANOVA). Percentages represent the variance in each parameter with age, sex and the interaction between age and sex.

**Supplementary Fig. 4:**
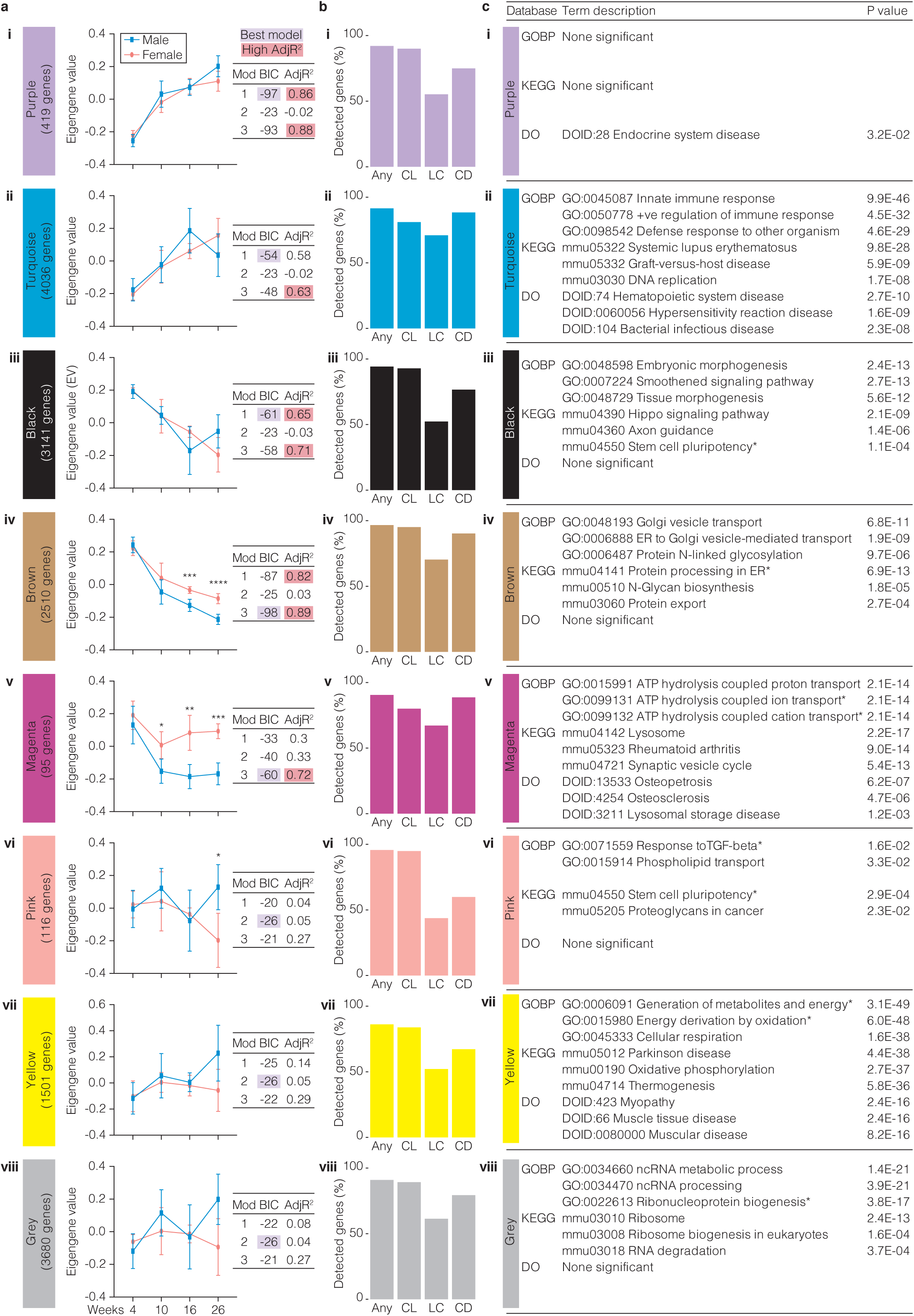
Gene clusters co-regulated during skeletal maturation. **a,** Pattern of gene expression change for each cluster (summarised by eigengenes) during postnatal skeletal maturation in male (blue) and female (red) mice **(i-viii)**. Significant separation of eigengene values between sexes at each age is indicated (* p<0.05, ** p<0.01, *** p<0.001). Three separate linear models fitting eigengene variance (EV) for each colour cluster (sex-only EV_cluster_∼Sex, age-only EV_cluster_∼Age, full model EV_cluster_∼Age + Sex + Age*Sex) to establish association between cluster expression age/sex are shown. The Bayesian Information Criterion (BIC) was used to select the best model fit and adjusted-R^2^ (AdjR^2^) used to estimate model strength. The optimum model (lowest value) selected by BIC is highlighted in purple. Linear models with a high AdjR^2^ (>0.6), indicating a strong association with eigengene expression, are highlighted in red. **b,** The percentage of expressed genes in each cluster that were also expressed in the osteocytic IDGSW3 cell-line (CL), in laser capture micro-dissected osteocytes (LC) and collagenase-digested bone samples (CD), or ANY of these orthogonal datasets. c, The top three most significantly enriched gene ontology biological processes (GOBP), KEGG pathways and disease ontology (DO) terms in each cluster ranked by p-value **(i-viii)** (Bonferroni corrected p<0.05).

**Supplementary Fig. 5:**
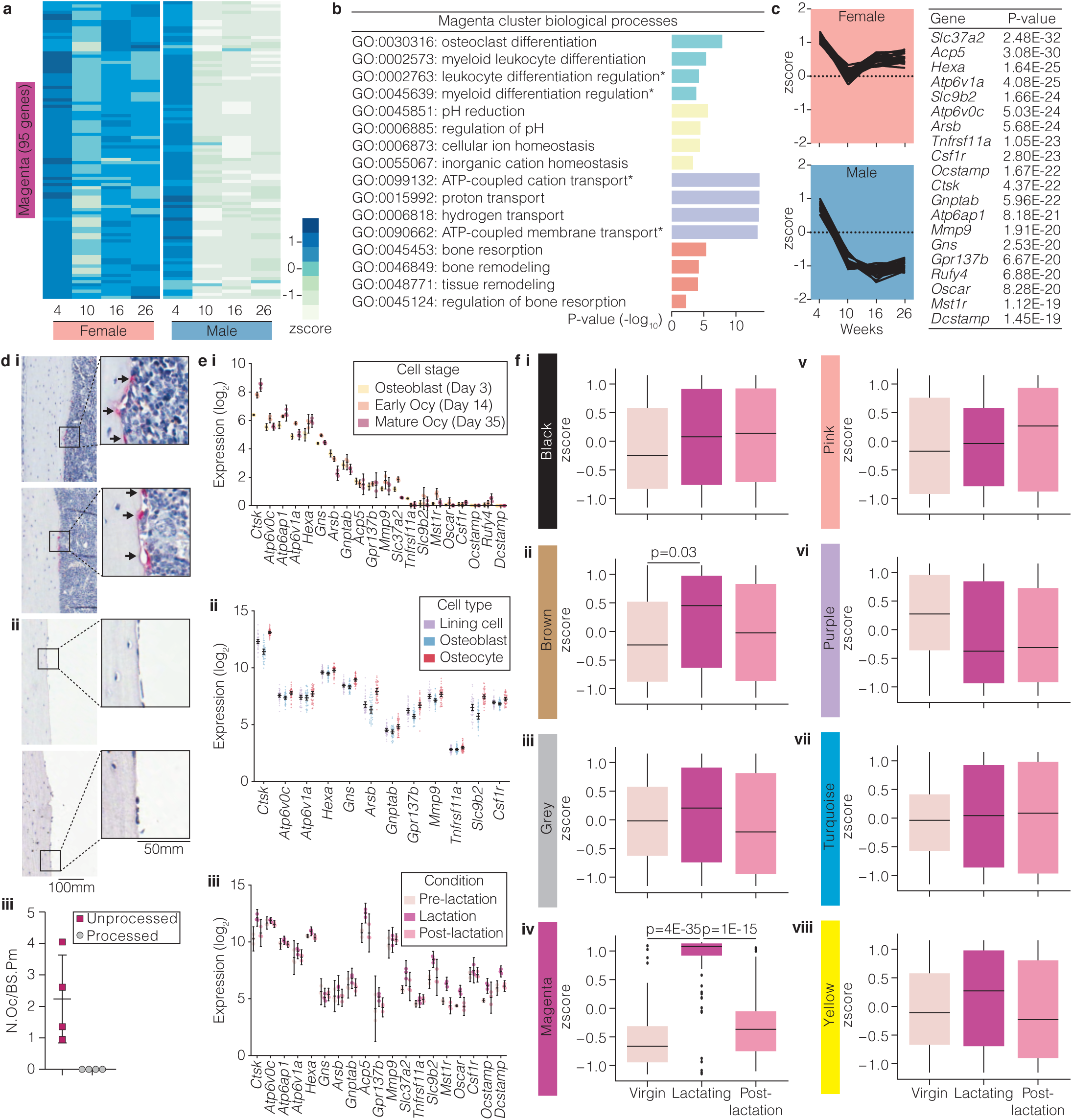
Magenta cluster genes associated with perilacunar-remodeling. **a,** Heatmap showing *Magenta* cluster gene expression at different ages (weeks) in male and female mice. Values reflect mean of scaled gene expression values at each time point. **b,** Biological processes significantly overrepresented among *Magenta* cluster genes grouped by semantic similarity (4 groups denoted by 4 different bar colours, p<0.05, *truncated description). **c,** Top 20 *Magenta* cluster genes associated with skeletal maturation in male and female mice (ranked by p-value). **d,** TRAP-stained histological sections of whole-bone (unprocessed) (i) and osteocyte-enriched (processed) (ii) bone samples and quantification of TRAP-positive osteoclasts on the bone surface (indicated by arrows) in each sample type (iii). **e,** Expression of the top 20 magenta genes in the osteocytic IDG-SW3 cell line (i), laser capture micro-dissected bone cells (ii) and collagenase digested osteocytes (iii). **f,** Expression of genes associated with each cluster in osteocytes from virgin mice, lactating mice and mice post-lactation (Post-lac)^15^. Tukey box-plots show a summary of median cluster gene expression values in each condition. Boxes indicate median and interquartile range (IQR) of scaled, normalized gene expression values, whiskers denote values ±1.5*IQR and outlier values beyond this range are shown as individual points. P-values were calculated by CAMERA^105^.

**Supplementary Fig. 6:**
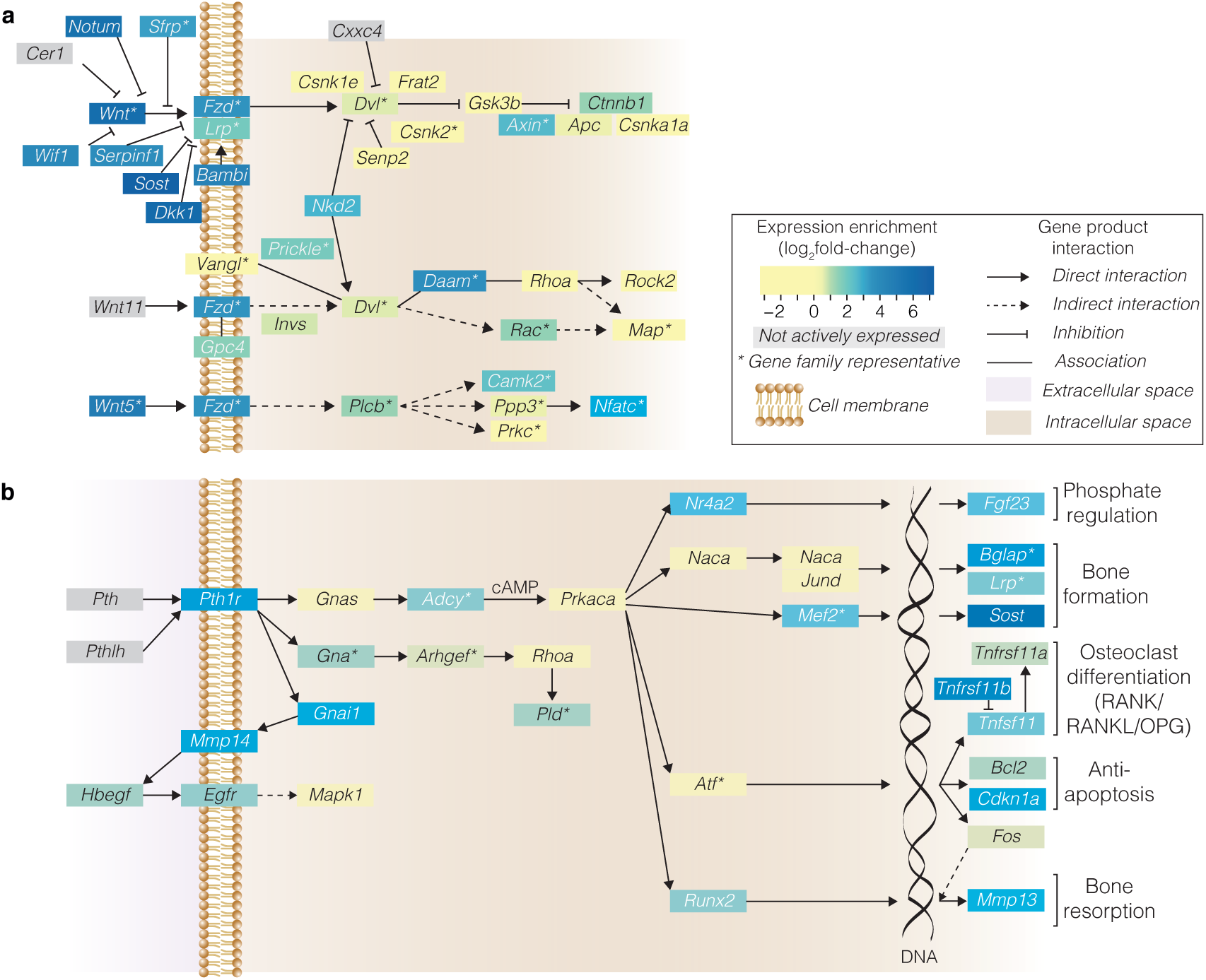
Signaling pathways with established roles in bone are enriched for expression in osteocytes. **a,** Wnt-signaling and **b,** PTH-signaling pathway genes actively expressed in osteocytes. Level of expression enrichment in osteocytes increases from yellow to blue (log_2_-scale). Grey represents genes not actively expressed. Lines and arrows indicate associations, inhibition, and direct or indirect interactions. * denotes most enriched gene where more than one molecule from a family of related genes can function at the same point in the pathway.

**Supplementary Fig. 7:**
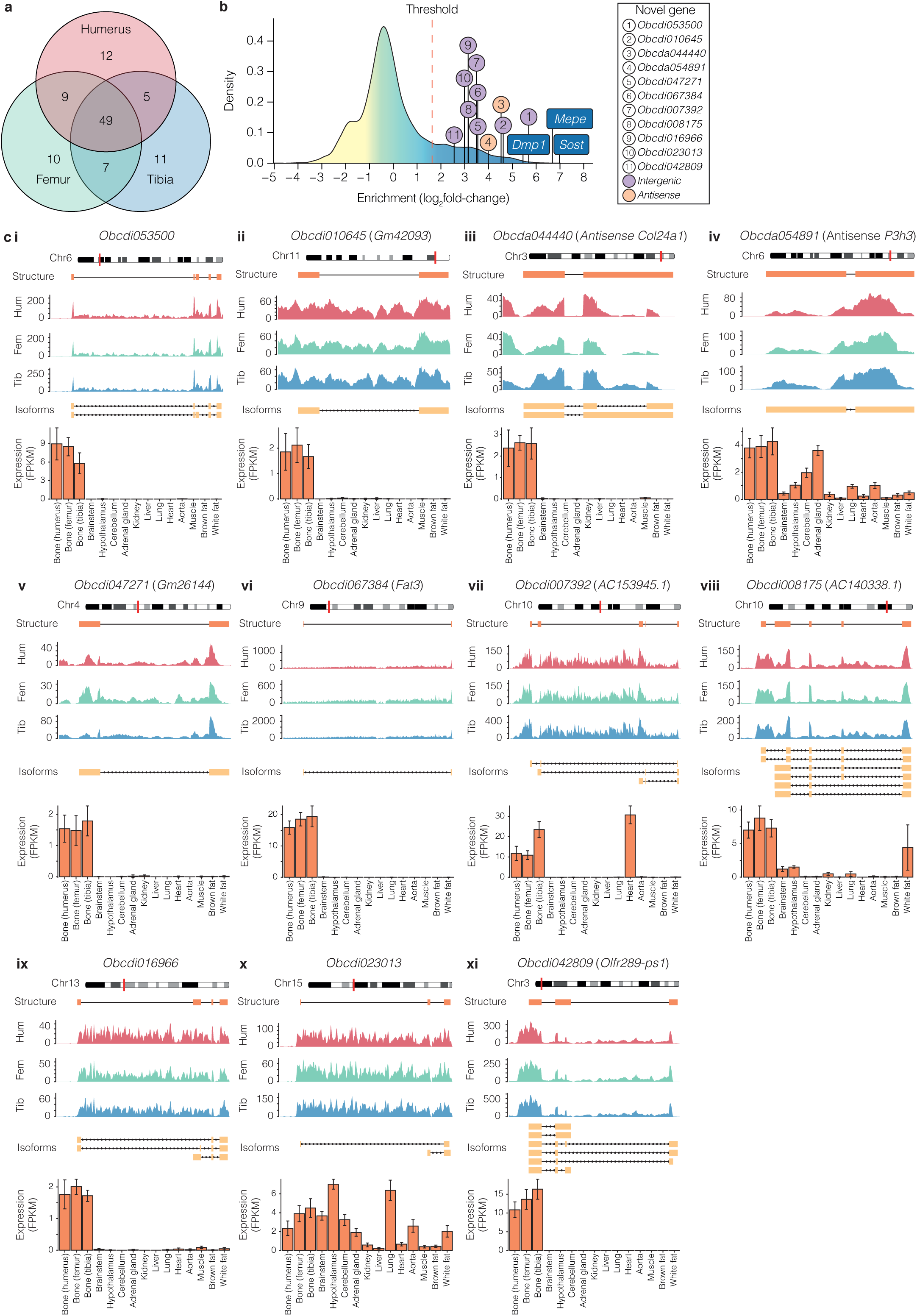
Novel genes identified in the *osteocyte transcriptome signature*. **a,** Venn diagram showing the number of novel genes actively expressed in osteocytes from different bone types. **b,** Expression enrichment of 11 novel genes identified in the *osteocyte transcriptome signature*. Intergenic genes are denoted in purple and antisense genes in orange. Established osteocyte genes are labeled with gene symbols. **c,** Gene structure diagrams of novel genes in the *osteocyte transcriptome signature* **(i-xi)**. Chromosome and location on each chromosome (red line), pooled read data alignment for humeri (Hum), femora (Fem) and tibiae (Tib) and individual predicted isoforms are shown. Histograms show normalized expression (mean FPKM ±SD) of each gene in osteocytes from three bones relative to 12 organs and tissues^28^.

**Supplementary Fig. 8:**
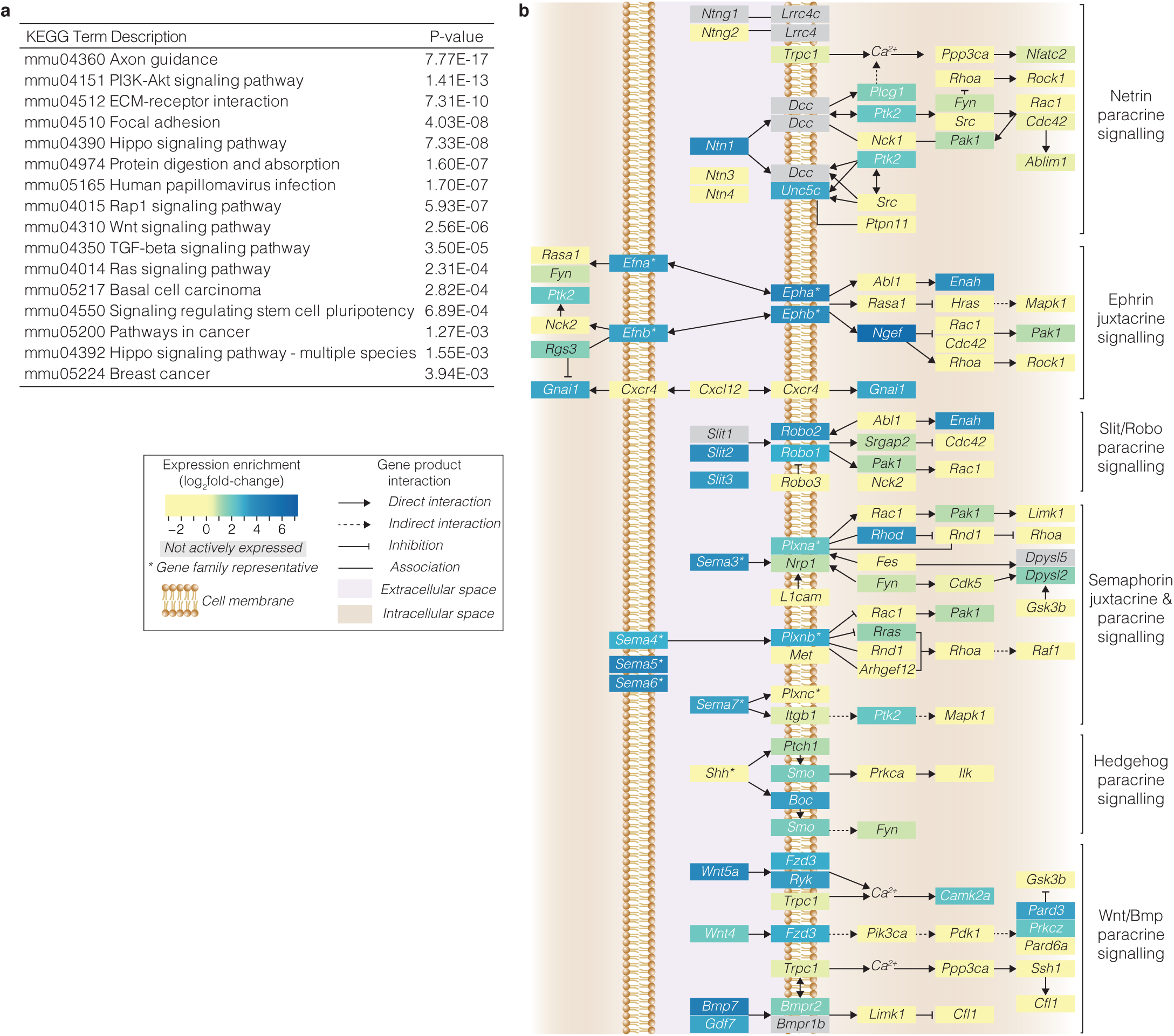
Axon guidance pathway genes are enriched in the *osteocyte transcriptome signature*. **a,** Top ranked KEGG pathways significantly enriched among *osteocyte transcriptome signature* genes and ranked by p-value (Bonferroni-corrected p<0.05). **b,** Axon guidance pathway genes actively expressed in osteocytes. Level of expression enrichment in osteocytes increases from yellow to blue (log_2_-scale). Grey represents genes not actively expressed. Lines and arrows indicate associations, inhibition, and direct or indirect interactions. * denotes most enriched gene where more than one molecule from a family of related genes can function at the same point in the pathway.

**Supplementary Fig. 9:**
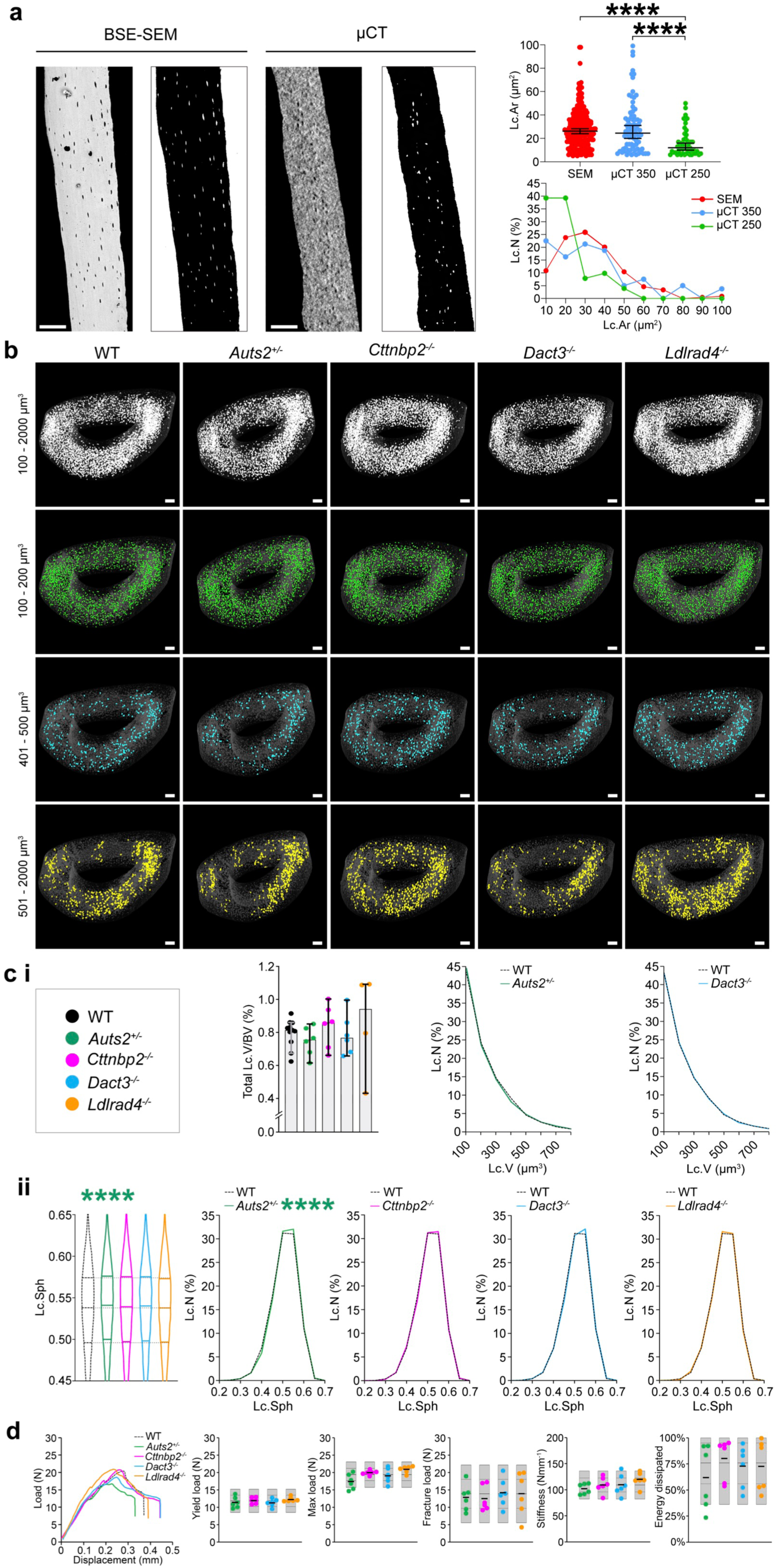
Disruption of the osteocyte network in mice with deletion of osteocyte signature genes. **a,** Longitudinal back-scattered electron scanning-electron microscopy (BSE-SEM) and micro-CT images of tibia cortical bone from adult female wild type (WT) mice. Micro-CT image generated at a 1µm voxel resolution using a segmentation threshold of 350 (equivalent to 852.72 mgHAcm^-3^). The corresponding images to the right show cortical bone in black and osteocyte lacunae in white. Graphs show the relationship between osteocyte lacuna area (Lc.Ar) and lacunar number (Lc.N) in BSE-SEM images and micro-CT images generated using a segmentation threshold of either 350 or 250 (equivalent to 552.5 mgHAcm^-3^). **** *P*<0.0001. **b,** Mid-tibia micro-CT images from adult female WT, *Auts2*^+/^, *Cttnbp2*^-/-^, *Dact3*^-/-^ and *Ldlrad4*^-/-^ mice showing distribution of osteocyte lacunae with volumes of 100-2000µm^3^, 100-200µm^3^, 401- 500µm^3^, and 501-2000µm^3^. Scale bar = 100µm. **c, (i)** Graphs show total osteocyte lacuna volume per bone volume (Lc.V/BV) in adult female WT, *Auts2*^+/-^, *Cttnbp2*^-/-^, *Dact3*^-/-^ and *Ldlrad4*^- /-^ mice and the distribution of Lc.V in *Auts2*^+/-^ and *Dact3*^-/-^ mice compared to WT (n = 4-11 per genotype). **(ii)** Violin plots and relative frequency graphs show distribution of osteocyte lacuna sphericity (Lc.Sph) in the four knockout mouse lines compared to WT (n = 4-11 per genotype). **** P<0.0001. **d,** Load displacement curves from femur 3-point bend testing of adult female WT, *Auts2*^+/-^, *Cttnbp2*^-/-^, *Dact3*^-/-^ and *Ldlrad4*^-/-^ mice. Dot plots show yield load, maximum and fracture loads, stiffness and energy dissipated prior to fracture. For each variable the mean (solid centre lines), ±1.0 SD (dotted lines) and ±2.0 SD (grey boxes) for WT mice (n=320) are shown. Mean values for each parameter in *Auts2*^+/-^, *Cttnbp2*^-/-^, *Dact3*^-/-^ and *Ldlrad4*^-/-^ lines are shown as a thick black line and individual data points as green, purple, blue and orange dots respectively (n=6 animals per genotype).

**Supplementary Fig. 10:**
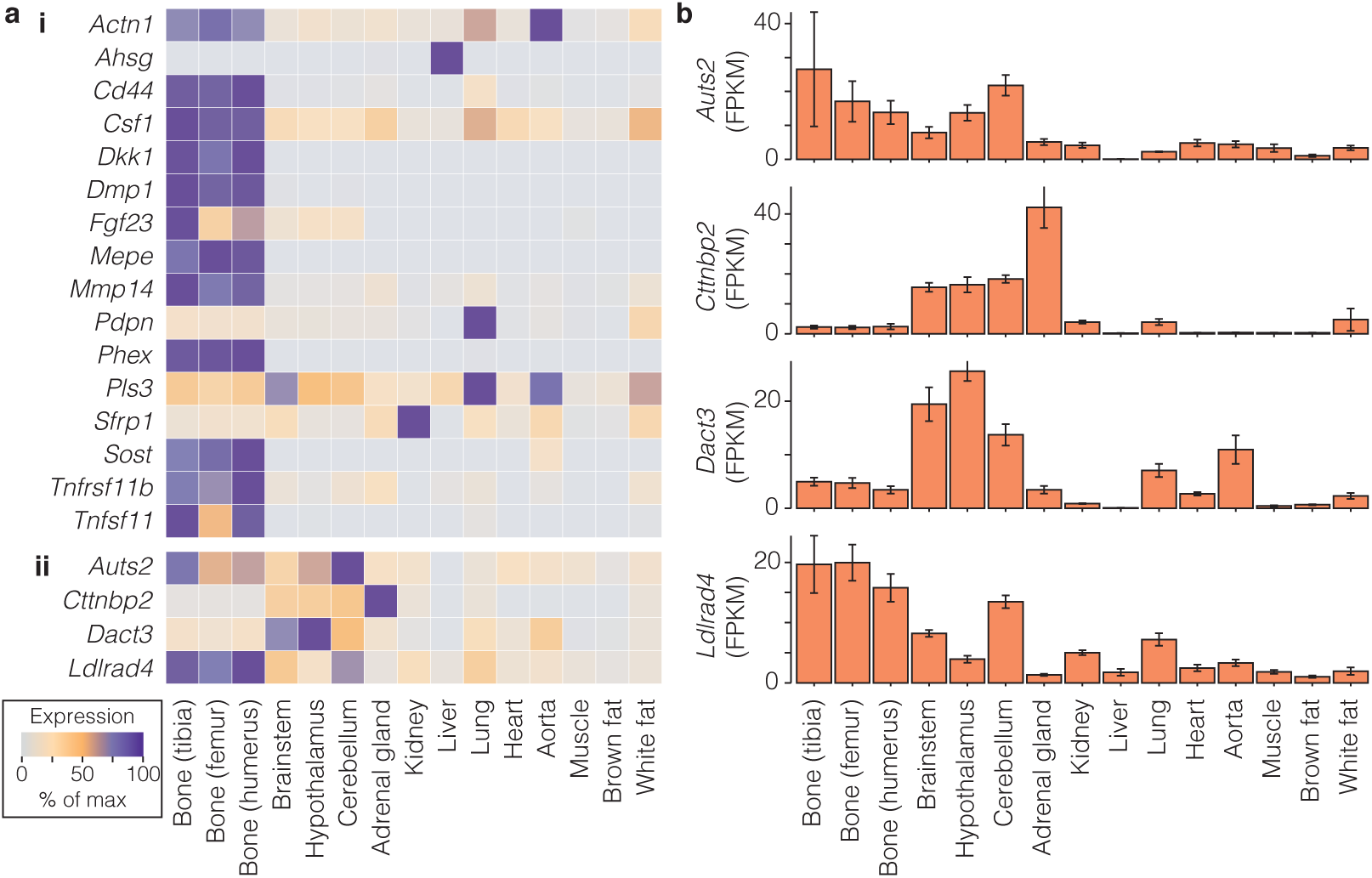
Expression of osteocyte signature genes associated with significant skeletal phenotypes across other organs and tissues. **a,** Relative expression in osteocytes and 12 non-skeletal tissues (% of maximum expression) of genes with established roles in the osteocyte network **(i)**, and four osteocyte transcriptome signature genes associated with significant skeletal phenotypes in the Origins of Bone and Cartilage Disease Program database **(ii)**. **b,** Normalised expression values (FPKM) of four osteocyte transcriptome signature genes associated with significant skeletal phenotypes in osteocyte-enriched bone and 12 non-skeletal tissues.

**Supplementary Fig. 11:**
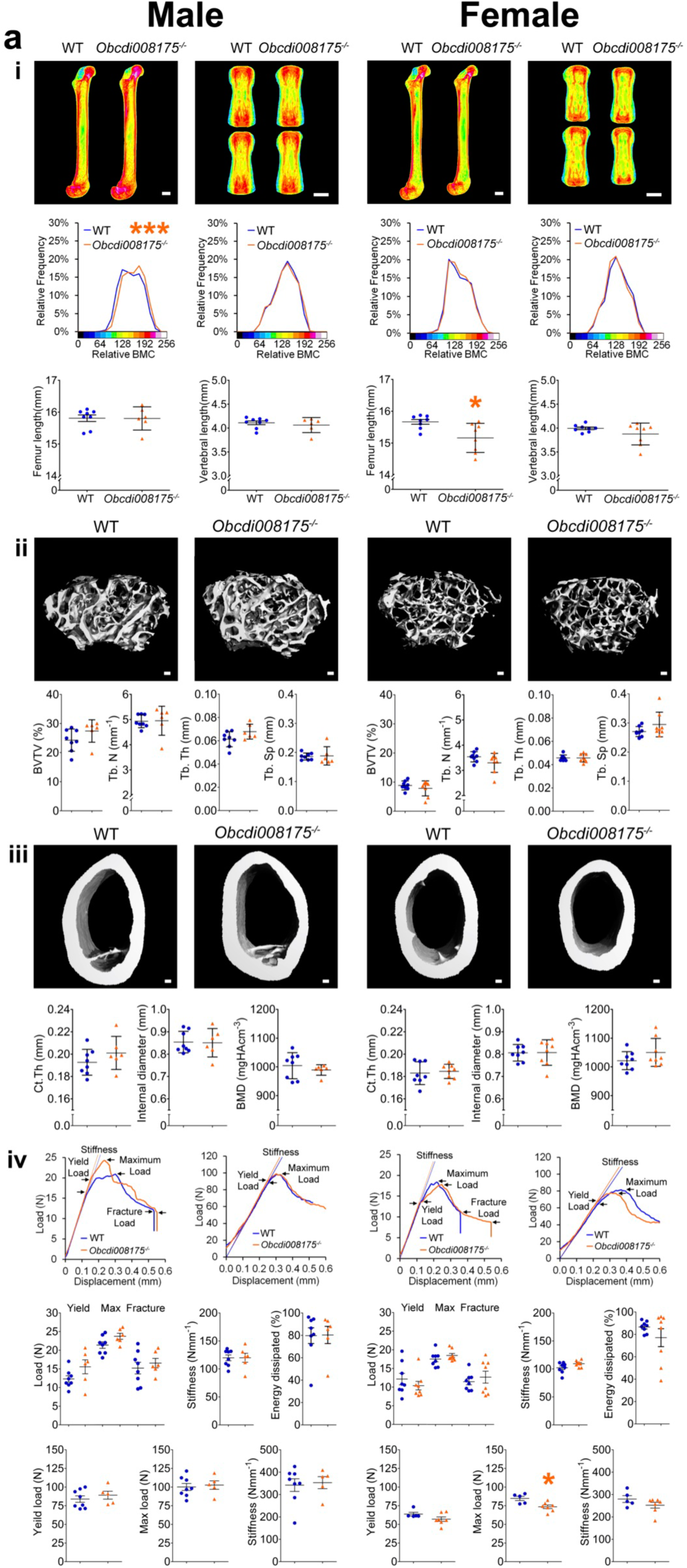
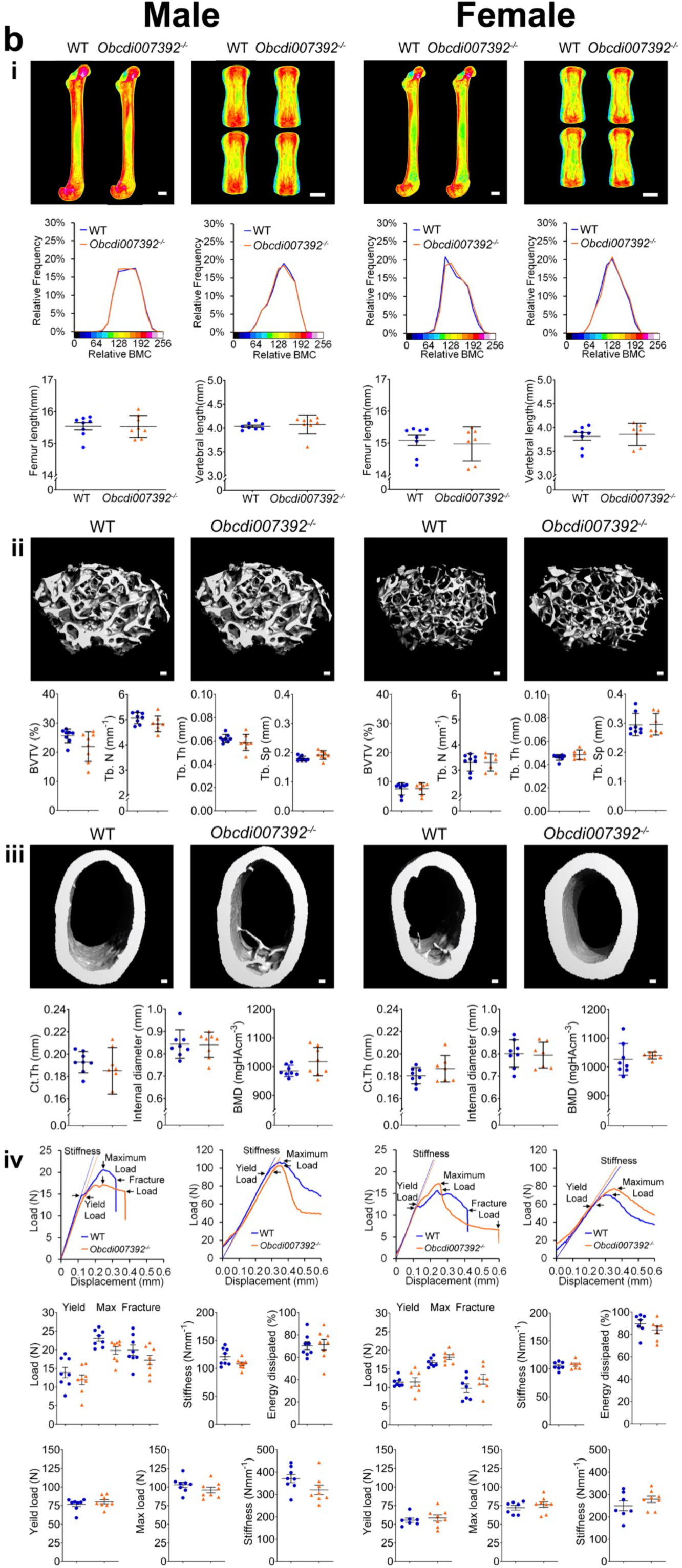
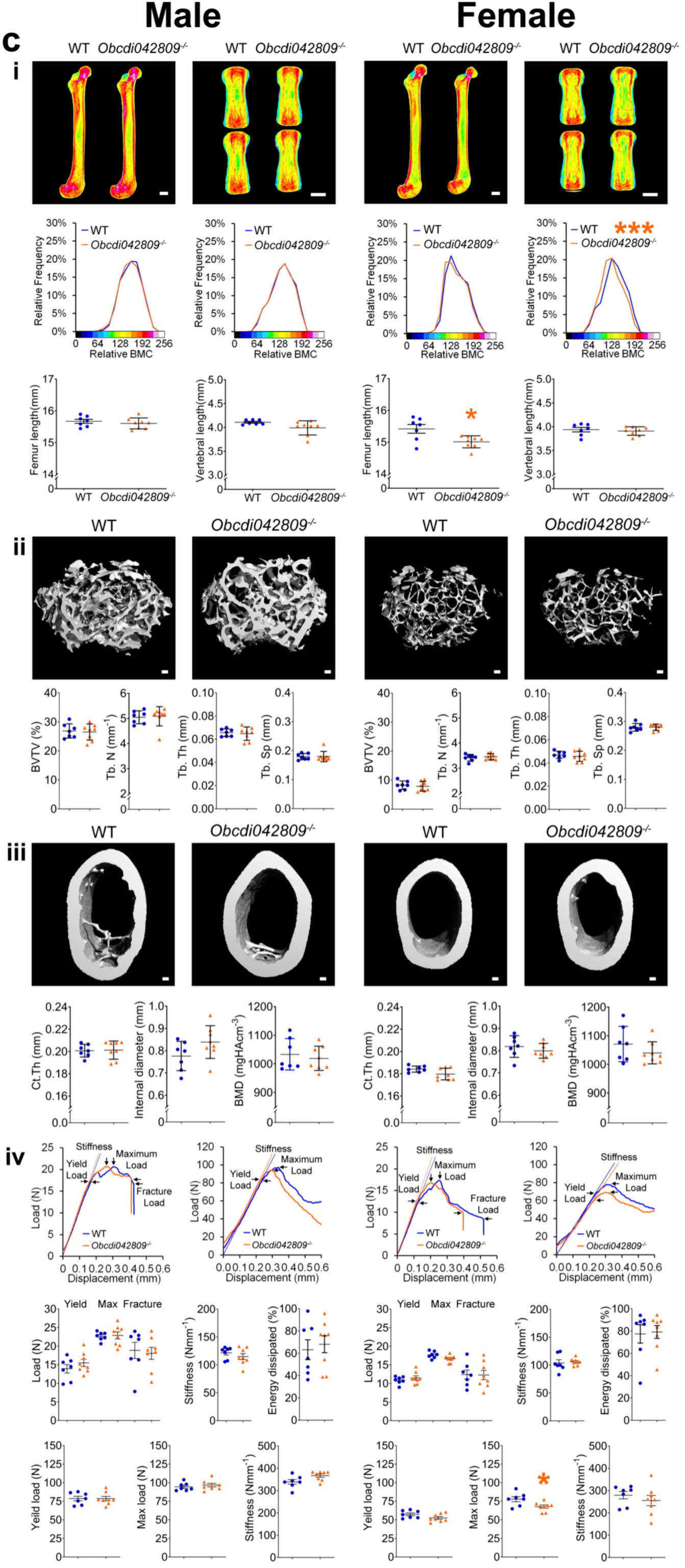
Deletion of novel *osteocyte transcriptome signature* genes effects skeletal structure and function. Skeletal phenotype of 16-week-old adult mice with deletion of novel genes identified in the osteocyte transcriptome signature. **a,** Mouse line *Obcdi008175^-/-^*, **b,** Mouse line *Obcdi007392^-/-^* and **c,** Mouse line *Obcdi042809^-/-^*. For each line, data from male mice are shown on the left and data from female mice on the right. Data from wild type animals are shown in blue and data from homozygous knockouts in orange. **(i)** Representative quantitative X-ray microradiographic images from the femurs and vertebrae. Pseudo-coloured images represent grey scale images using a 16-colour interval scheme with low mineral content blue and high mineral content red. Scale barL=L1Lmm. Top graphs show relative frequency histograms of femur bone mineral content (BMC) left and vertebra BMC right (n=6–8 per sex, per genotype, ***P<0.001 versus WT; Kolmogorov-Smirnov test). Bottom graphs show femur lengths and caudal vertebral heights (mean ± SD, n=6–8 per sex, per genotype, *P<0.05, versus WT; unpaired *t*-test). **(ii)** Representative micro-CT images of distal femur trabecular bone. Scale barL=L100µm. Graphs show bone volume as a proportion of tissue volume (BV/TV), trabecular number (Tb.N), trabecular thickness (Tb.Th), trabecular separation (Tb.Sp) (mean ± SD n=6-8 per sex, per L 100µm. Graphs show cortical thickness (Ct.Th), internal endosteal diameter and bone mineral density (BMD) (mean ± SD n=6-8 per sex, per genotype). **(iv)** Representative load displacement curves from 3-point bend testing of the femur (left) and caudal vertebrae compression testing (right). Black arrows indicate yield, maximum and fracture loads and stiffness is indicated by straight blue (WT) or orange (knockout) lines. Top graphs show yield load, maximum and fracture loads, stiffness and energy dissipated prior to fracture (toughness) from femur 3-point bend testing (mean ± SD, n=6–8 per sex, per genotype). Bottom graphs show yield and maximum loads and stiffness from caudal vertebrae compression testing (mean ± SD, n=6–8 per sex, per genotype, *P<0.05, versus WT; unpaired t-test).

**Supplementary Fig. 12:**
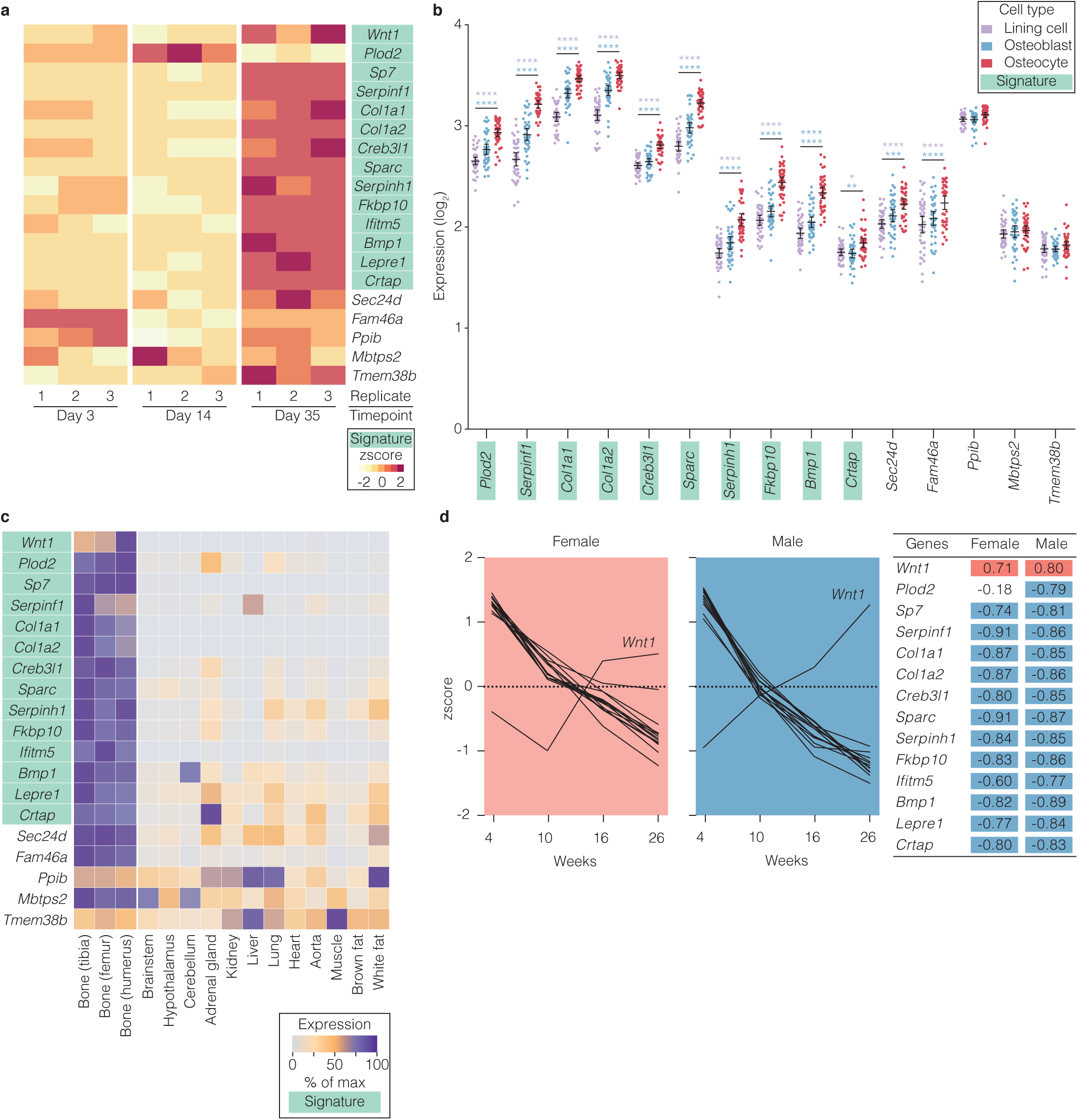
Expression of osteogenesis imperfecta genes in the *osteocyte transcriptome signature*. **a,** Expression of osteogenesis imperfecta (OI) genes during osteocyte differentiation from osteoblast-like cells (day 3) to early (day 14) and late osteocytes (day 35) (z-score of normalized counts)^50^. Numbers represent replicates. *Osteocyte transcriptome signature* genes are highlighted in green. **b,** Expression of OI-genes in rat osteocytes (red), osteoblasts (blue) and bone lining cells (purple) isolated by laser capture micro-dissection^16^. 15/19 OI-genes had rat-orthologs on the original microarray and all are shown. Dots represent individual sample expression values (n=40 per cell type). *Osteocyte transcriptome signature* genes are highlighted in green. Significant differences in expression are indicated by ** (P<0.01), *** (P<0.001) and **** (P<0.0001), with colours denoting comparisons between osteocyte and osteoblasts (blue) and between osteocytes and bone lining cells (purple). **c,** Heatmap showing expression of OI-genes in osteocytes isolated from the tibia, femur and humerus relative to 12 organs and tissues^28^ (shown as percentage of maximum mean-FPKM). *Osteocyte transcriptome signature* genes are highlighted in green. **d,** OI-gene expression in osteocytes with age (4-26 weeks) in female (pink) and male (blue) mice during skeletal maturation. Lines represent individual OI-genes. Pearson correlations between genes and age are tabulated. Genes significantly correlated with age are highlighted (negative = blue, positive = red, p<0.05).

## Supplementary Tables

**Supplementary Table 1.** Sample specific gene activity thresholds

**Supplementary Table 2.** The osteocyte-enriched transcriptome

**Supplementary Table 3.** Osteocyte transcriptome genes differentially expressed between bones

**Supplementary Table 4.** Skeletal maturation clusters

**Supplementary Table 5.** The *osteocyte transcriptome signature*

**Supplementary Table 6.** Gene ontology terms enriched in the *osteocyte transcriptome signature*

**Supplementary Table 7.** Novel genes expressed in osteocytes

**Supplementary Table 8.** *Osteocyte transcriptome signature* genes deleted in mice and phenotyped by the origins of bone and cartilage disease program

**Supplementary Table 9.** Expression of genetic skeletal disorder genes in the osteocyte network

**Supplementary Table 10.** *Osteocyte transcriptome signature* genes associated with eBMD in the UK Biobank cohort

**Supplementary Table 11.** *Osteocyte transcriptome signature* genes associated with OA in the UK Biobank cohort

